# Ready-to-use nanopore platform for the detection of any DNA/RNA oligo at attomole range using an Osmium tagged complementary probe

**DOI:** 10.1101/2020.10.05.327460

**Authors:** Albert S. W. Kang, Janette G. Bernasconi, William Jack, Anastassia Kanavarioti

**Author notes:** Correspondence should be addressed to A.K.

## Abstract

Nanopores can serve as single molecule sensors. We exploited the MinION, a portable nanopore device from Oxford Nanopore Technologies (ONT), and repurposed it to detect any DNA/RNA oligo (target) in a complex mixture by conducting voltage-driven ion-channel measurements. The detection and quantitation of the target is enabled by the use of a unique complementary probe. Using a validated labeling technology, probes are tagged with a bulky Osmium tag (Osmium tetroxide 2,2’-bipyridine), in a way that preserves strong hybridization between probe and target. Intact oligos traverse the MinION’s nanopore relatively quickly compared to the device’s acquisition rate, and exhibit count of events comparable to the baseline. Counts are reported by a publicly available software, *OsBp_detect*. Due to the presence of the bulky Osmium tag, probes traverse more slowly, produce multiple counts over the baseline, and are even detected at single digit attomole (amole) range. In the presence of the target the probe is “silenced”. Silencing is attributed to a 1:1 double stranded (ds) complex that doesn’t fit and can’t traverse this nanopore. This ready-to-use platform can be tailored as a diagnostic test to meet the requirements for point-of-care cell-free tumor DNA (ctDNA) and microRNA (miRNA) detection and quantitation in body fluids.

## INTRODUCTION

Blood draw or other body fluid samples, known as liquid biopsies^1-10^, contain pertinent information regarding the health status of an individual, the progress of a disease, whether or not an individual is disease-free after surgery, and even whether or not a certain therapy strategy seems promising^2^. Liquid biopsies are far less invasive procedures than surgical/tumor biopsy. Body fluids contain cell-free DNA, a small fraction of which originates from the tumor. In 2016 the US Food and Drug Administration approved the first liquid biopsy test for EGFR-activating mutations in patients with non-small-cell lung cancer as a companion diagnostic test to enable therapy selection^10^. Cell-free DNA is believed to be fragmented and shorter in diseased individuals compared to healthy^3^. In addition to DNA, body fluids contain long non-coding RNAs^11^, tRNA derived fragments^12^, and other RNA oligos in the range of 20 to 300 nucleotides (nt). Among them is a group of single-stranded (ss) RNAs, 17 to 25nt long, microRNAs or miRNAs^13-15^. They were discovered 20 years ago and proven to be the tiny regulators that control the post-transcriptional expression of proteins^16,17^; they are highly conserved and surprisingly stable in body fluids^13,18^. Currently there are over 2,300 human miRNAs known^19^, the subject of over 100,000 scientific publications. Up- or down-regulation of miRNAs is associated with various human diseases including cancer, heart disease, kidney disease, obesity, diabetes, etc.^17,20^. miRNAs are proposed as biomarkers^21,22^ and as potential therapeutics^23^ in personalized medicine. Body fluids contain trace amounts of ctDNA and miRNAs, that require simple, validated, and highly sensitive assays for testing^8,9,24-27^. Current technologies for “small RNA” identification and quantitation, include microarray, NGS sequencing (Ion Torrent or Illumina (small RNA-seq)), and qRT-PCR-based methods that have been employed so far with great success^24-27^. These technologies require substantial infrastructure and skilled personnel, are not well-suited for point-of-care testing and are out of the question for home testing. With our technology, this information could be easily accessible, even in the privacy of one’s home.

The last 30 years have seen a surge in nanopore-based technologies using either solid-state or protein nanopores^28-34^. In 2014 Oxford Nanopore Technologies (ONT) introduced and commercialized the first portable nanopore device to sequence DNA and RNA practically anywhere, as long as a computer and internet are available^34^. ONT technology is based on the CsGg protein nanopore^35^, with sub 2nm diameter, inserted in a planar lipid bilayer membrane that separates two electrolyte filled compartments (Fig. 1a). Applying a voltage across the two compartments leads to a constant flow of electrolyte ions (*I*_*o*_) via the pore, recorded as a function of time (*i-t*). The passage of a single molecule through the pore reduces *I*_*o*_ to a lower level of residual ion current (*I*_*r*_). This is recorded as an “event” with (*I*_*r*_) and residence time (t) (Fig. 1b). Currently the ONT platform is exclusively used for DNA/RNA sequencing^34^, while comparable nanopore platforms are successfully employed for single molecule analyses^36-51^. These and other studies, using experimental platforms, introduced nanopores to the arena of minimally invasive diagnostic assays.

**Fig. 1:**
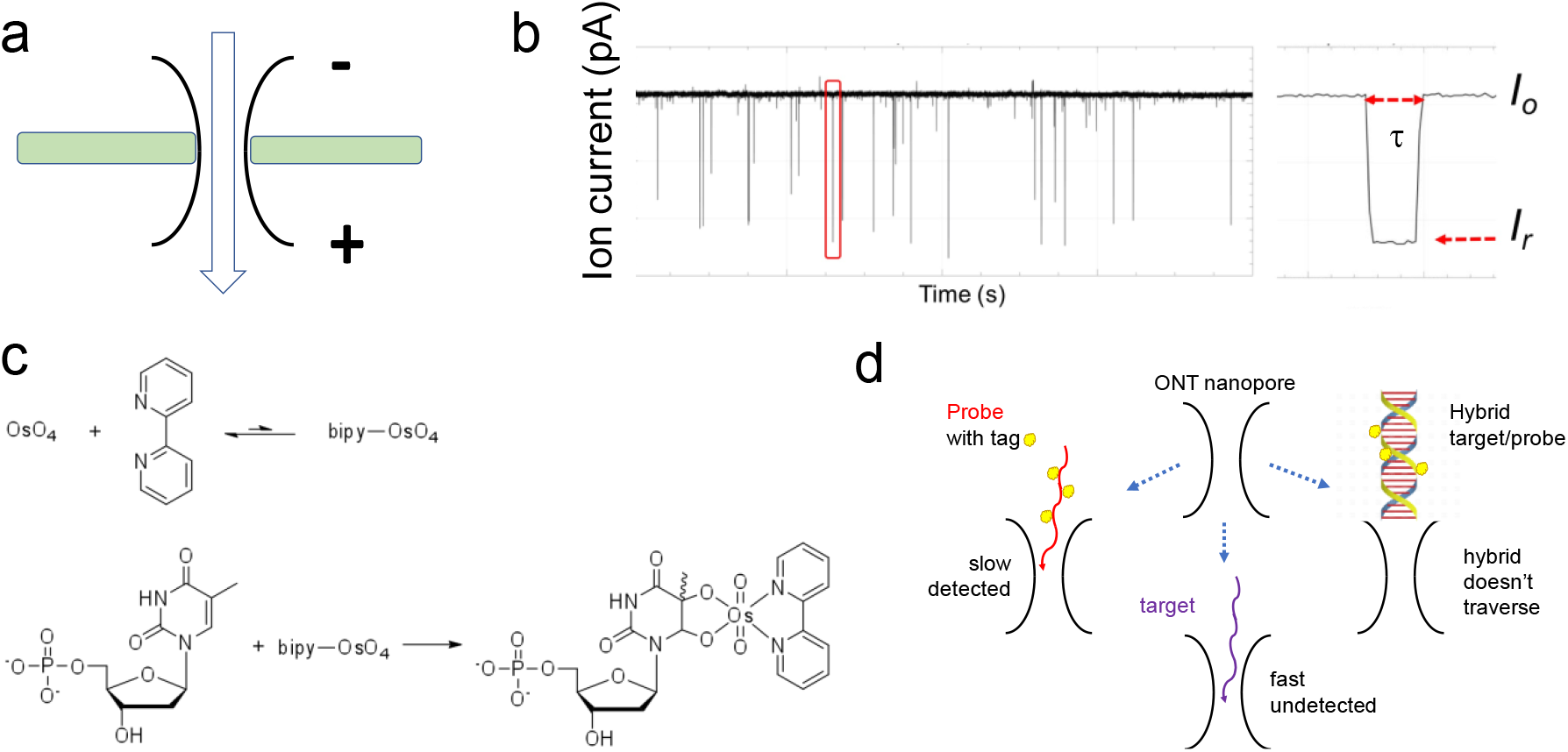
The critical parameters of the proposed diagnostic assay. **(a)** Schematic representation of a nanopore within a planar bilayer lipid membrane that separates two electrolyte filled compartments. Applying a constant voltage to the flow cell guides the passage of ions through the nanopore creating a measurable ionic current. **(b)** The *i-t* trace obtained from a voltage-driven ion-channel experiment where the constant flow of electrolyte ions (*I*_*o*_) via the pore is interrupted by the passage of molecules. These molecules appear as “events” with residual ion current *I*_*r*_ and residence time t. **(c)** OsBp labeling reaction: OsO_4_ and 2,2-bipyridine (bipy) have a low association constant, but their mixture adds to the C5-C6 double bond of pyrimidines and forms a stable conjugate, called here osmylated nucleic acid. The addition of OsBp creates a chromophore that absorbs in the range of 312 nm where native nucleic acids do not absorb (see Experimental Section). **(d)** Cartoon illustrates the concept behind the proposed diagnostic test. ssDNA and ssRNA traverse the MinION’s CsGg nanopore and exhibit few counts because they traverse faster, compared to the device’s relatively slow acquisition rate; ds nucleic acids are too big and do not traverse this nanopore. Despite being bulkier than ss native nucleic acids, osmylated ss nucleic acids traverse the MinION/CsGg pore and produce numerous events. When an osmylated nucleic acid (probe) is added to a sample that contains its complementary (target), the probe and the target form a hybrid. When the target’s concentration is equal or higher than the probe’s concentration, the probe is hybridized. The probe is then prevented from traversing the pore, and few or no events are observed. The absence of target in the sample is evidenced by the numerous events produced by the probe while it freely traverses the pore.

The appeal of using nanopores as single molecule sensors resides in their ability to detect and count every molecule that traverses the pore. To appreciate this advantage, let us consider a 10mL sample that contains miRNA21 at 10×10^−15^M (10 femtoM or 10 fM) concentration. To the best of our knowledge, there is no method to analyze such diluted sample without amplification. This sample contains 1×10^−19^ moles, i.e. 0.1 attomole (amole), or approximately 60,000 molecules of miRNA21. A size-suitable nanopore may detect some, if not all, as long as there is an electrical field to guide them towards the pore and thread them through it. Compared to other analytical tools, nanopores appear highly fitting for trace analysis such as the DNA/RNA oligos in body fluids. The argument becomes even more compelling when there is a ready-to-use microfluidic device with an array of about 500 actively reporting nanopores which is the MinION configuration from ONT^34^.

ONT commercialized portable nanopore devices that carry two types of flow cells; the MinION with 512 channels and the Flongle with 126 channels, all monitored simultaneously^34^. ONT promotes these devices for direct sequencing of DNA and RNA with a minimum length of 200 nucleotides (nt)^34^. In a recent study, attempts to sequence RNAs shorter than 200nt following ONT protocol were unsuccessful^52^. In another study 100nt deoxyoligos were successfully sequenced by the MinION, albeit only after circularization and rolling circle amplification to produce long DNA with 100nt repeats^53^. Most of the DNA/RNA available in biological fluids is fragmented with lengths estimated in the range of 200bp^3^ and miRNAs are very short^16^, and not amenable to direct ONT sequencing protocols. Still ONT devices/flow cells can be used for single molecule ion-channel measurements, just like any other nanopore. While several experimental nanopore platforms were successfully used for miRNA profiling (see later), we believe to be the first to show feasibility using a commercial device^49^. The advantage compared to other platforms is that the ONT technology is accessible, ready-to-use, relatively inexpensive, and does not require any special skills or infrastructure.

We have exploited selective labeling of nucleic acids in an effort to enhance base-to-base discrimination^41,44,49,54^, and used Osmium tetroxide 2,2’-bipyridine (OsBp) as the label/tag. OsBp is not reactive towards the purines and does not cleave the phosphodiester bond in DNA or RNA. OsBp adds to the C5-C6 double bond of the pyrimidines and forms two strong C-O bonds without cleaving the pyrimidine ring (Fig. 1c)^55-60^. The products are named here “osmylated” nucleic acids. Reactivity of OsBp towards thymidine (T) is 28- and 7.5-times higher compared to the reactivity towards deoxycytidine (C) and deoxyuridine (U), respectively^44^. We demonstrated control over the labeling conditions, and developed a protocol to selectively label T in the presence of the other pyrimidines^58^. We developed capillary electrophoresis (CE) and High-performance Liquid chromatography (HPLC) methods to measure the extent of labeling in short and long DNA and RNA^58,59^ (Experimental Section). We also conducted voltage-driven ion-channel measurements using SiN solid-state nanopores^41^, the α-Hemolysin nanopore^44^, as well as the CsGg proprietary nanopore in the MinION^49^, and demonstrated that all three platforms allow the translocation of osmylated nucleic acids, and clearly discriminate them from native nucleic acids. The discrimination is manifested as an event with markedly lower *I*_*r*_ and longer t with increasing number of OsBp moieties (Fig. 1b). Here we exploit these features in order to single-out, detect, and count OsBp-tagged oligos in a complex mixture of native DNA and RNA.

Fig. 1d illustrates the concept behind nanopore-based identification and quantification of a target oligo in a complex mixture; it is enabled by a custom-designed OsBp-tagged deoxyoligo (probe), that is complementary to the target. The use of a complementary oligo as a probe has been validated in several experimental nanopore platforms^38,42,43,45,51^. These platforms however are made complicated by the fact that the probe is conjugated to a protein^38^, a nanoparticle^51^, a homopolymer^42^, or a polypeptide^45^. The detection in these platforms relies on counting the long blockades produced by the hybrid bumping at the nanopore entry and practically “getting stuck”. None of these methods reached commercial availability which hinders their broader use. In contrast to earlier approaches based on detection of the hybrid^38,42,43,45,51^, our technology detects the translocation of the bulky osmylated probe facilitated by the relatively slow acquisition rate of the MinION (3.012 kHz sampling rate, equivalent to reporting 3 data points per 1 ms)^34^. While the slow sampling rate misses most translocation events of native DNA/RNA oligos, the events corresponding to the translocation of our tagged probes are detected. In the absence of the target, the probe traverses the nanopore and produces a detectable event. In the presence of the target, the probe forms a 1:1 ds complex (hybrid) with the target. The hybrid doesn’t fit and can’t traverse the proprietary CsGg nanopore of the ONT devices^34^ (Fig. 1d). Hence the probe is “silenced”. The hybrid doesn’t prevent ss nucleic acids going through the pore, as ONT technology incorporates automatic reversal of voltage to free the nanopores from such unproductive occurrences. Therefore, to test the presence/absence of a target oligo in a sample, a probe is simply added to the sample and the mixture is tested in an ONT device by conducting voltage-driven single molecule experiments. Absence of the target oligo in a sample is conjectured from the observation of numerous events due to the probe’s translocation via the pore. The presence of the target oligo in a sample is concluded from the absence of events, due to hybrid formation between the probe and the target. Quantification of the target is based on the known concentration of the probe and the 1:1 hybrid formation. The concept is neither unique to the Osmium tag nor to the ONT nanopore array. Any bulky tag with selectivity for one of the bases should yield comparable behavior to what is observed with OsBp. In addition, most nanopore platforms with dimensions that allow the translocation of ss nucleic acids and prevent the translocation of ds nucleic acids will conform to the concept described in Fig. 1d, under suitable conditions of applied voltage and data acquisition speed.

## RESULTS AND DISCUSSION

The materials used in this study are all synthetic oligos of the highest purity and are listed in the table. Osmylation protocols were developed by us. Intact and osmylated nucleic acids were further characterized in-house by validated HPLC methods^59,61^ (see Experimental Section). Nanopore experiments were conducted using the ONT devices and the ONT supplied Flush buffer (ONT buffer or buffer), in addition to company’s instructions of how to prime the flow cell, add the sample, select voltage, and acquire the raw *i-t* traces. No sample library was prepared and no enzyme-assistance was exploited. Samples were prepared in 90-95% ONT buffer, unless otherwise noted. The nanopore experiments reported here were conducted at the factory-preset, flow cell temperature in the range of 34-35°C. The raw *i-t* traces of all channels (a *fast5* file) were captured and analyzed using *OsBp_detect* software^62^ (see Experimental Section). The output, a *tsv* file, is read using Microsoft Excel. It lists *I*_*0*_ value for each channel, as well as the selected events and their *I*_*r*_ value from which *I*_*r*_/*I*_*o*_ is calculated. It also lists the times, in data time points, of the beginning and of the end of each event (see Fig. S1, Supplementary Information). *OsBp_detect* permits manual setting of parameters, in order to select events of interest^62^. Here we selected events with residence time 4 ≤ t ≤ 300 data time points or the equivalent 1.3≤ t ≤ 100ms, and fractional residual ion current *I*_*r*_/*I*_*o*_ ≤ 0.55 (Fig. 1b). Figs 2, 4 through 7 present histograms of count of events (abbreviated as counts or events) as a function of *I*_*r*_/*I*_*o*_ using a 0.05 bin.

**Fig. 2:**
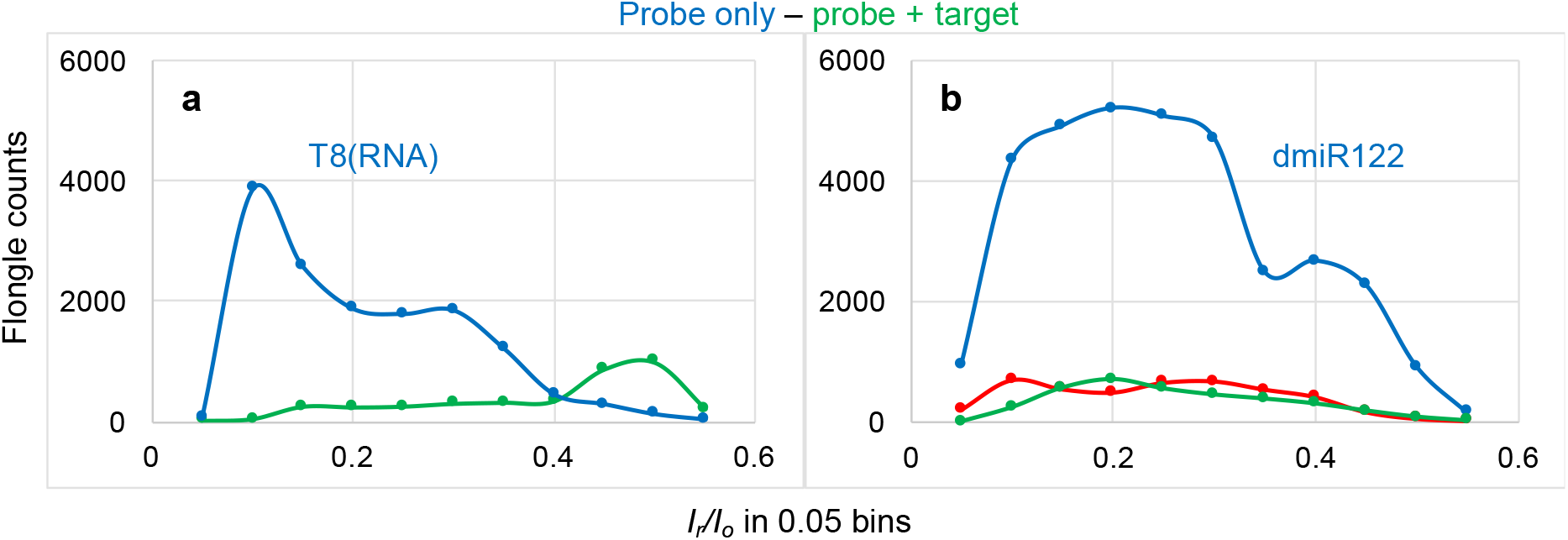
Voltage-driven ion-channel (nanopore) experiments conducted with the Flongle ONT device; samples in > 90% ONT buffer. **(a)** 1h at −200mV using the same Flongle flow cell: (i) 5μM probe T8(RNA) (blue trace) and (ii) 5μM each a mixture of T8(RNA) and d(CT)_10_ (green trace). It is noticeable that these two molecules are only partially complementary to each other. Count of events (counts) were obtained using the *OsBp_detect* software to analyze and report the raw *fast-5* file data acquired with MINKNOW (see Experimental Section). Counts are plotted as a function of *I*_*r*_/*I*_*o*_ with a bin size of 0.05. **(b)** 2h at −190mV using the same Flongle flow cell: (i) 5μM probe dmiR122 (blue trace), (ii) 5μM each a mixture of dmiR122 and miRNA122 (green trace), and (iii) 1h at −180mV a mixture 10μM each of miRNA122 and miRNA140 (red trace) using a different Flongle flow cell. It is noticeable that dmiR122 carries 4 OsBp moieties and its sequence is perfectly complementary to miRNA122. Data acquisition and analysis as described under (a).

**Fig. 3:**
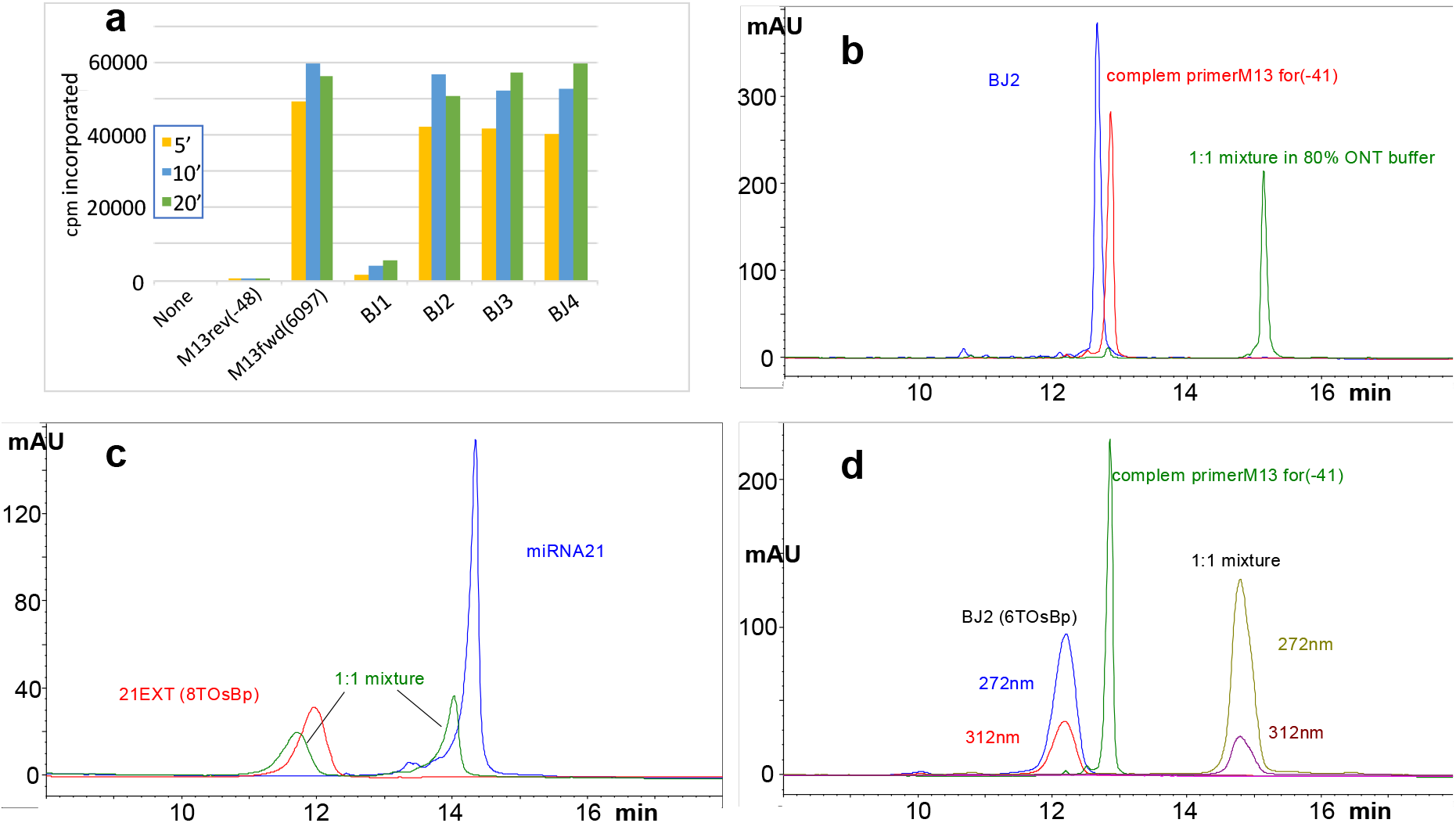
Alternative approaches to testing hybridization between osmylated probes and targets. **(a)** Enzymatic elongation of osmylated primers using ssM13mp18 DNA as the template and DNA polymerase; time points obtained at 5, 10 and 20min. No primer and M13*rev*(−48) used as negative controls. With the exception of BJ1, all the other osmylated primers exhibit enzymatic elongation comparable to the positive control M13*fwd*(6097). Absence of elongation with BJ1 is attributed to the presence of a T(OsBp) at the 3’end. **(b), (c) and (d)** Overlapping HPLC profiles from the analyses of different samples, with samples at about 5μM in about 90% ONT buffer. The same HPLC method B was used for all the samples (see Experimental Section). Intact ss oligos and ds oligos appear as sharp peaks, whereas osmylated oligos and hybrids with one osmylated strand appear as broad peaks; hybrids elute later compared to ss nucleic acids. **(b)** Sample composition: intact BJ2 (blue trace), intact complement of primerM13*for*(−41) (red trace). The HPLC profile of their equimolar mixture is consistent with hybridization (green trace). **(c)** Sample composition: miRNA21 (blue trace), probe 21EXT carrying 8 dT(OsBp) moieties (red trace), and equimolar mixture of the two (green trace). HPLC profile of the mixture consistent with NO hybridization, attributed to the high number of single OsBp tags, 6 within a sequence of 22nt, likely to distort the helical structure of the probe, and prevent ds formation. **(d)** Sample composition practically the same as under (b), with the exception that in these samples BJ2 carries 6 dT(OsBp) moieties (first peak with broad shape and absorbance at 312nm). HPLC profile of the mixture is consistent with hybridization, attributed to the fact that most of the OsBp moieties are adjacent, so that the rest of the sequence can still hybridize with the target.

**Fig. 4:**
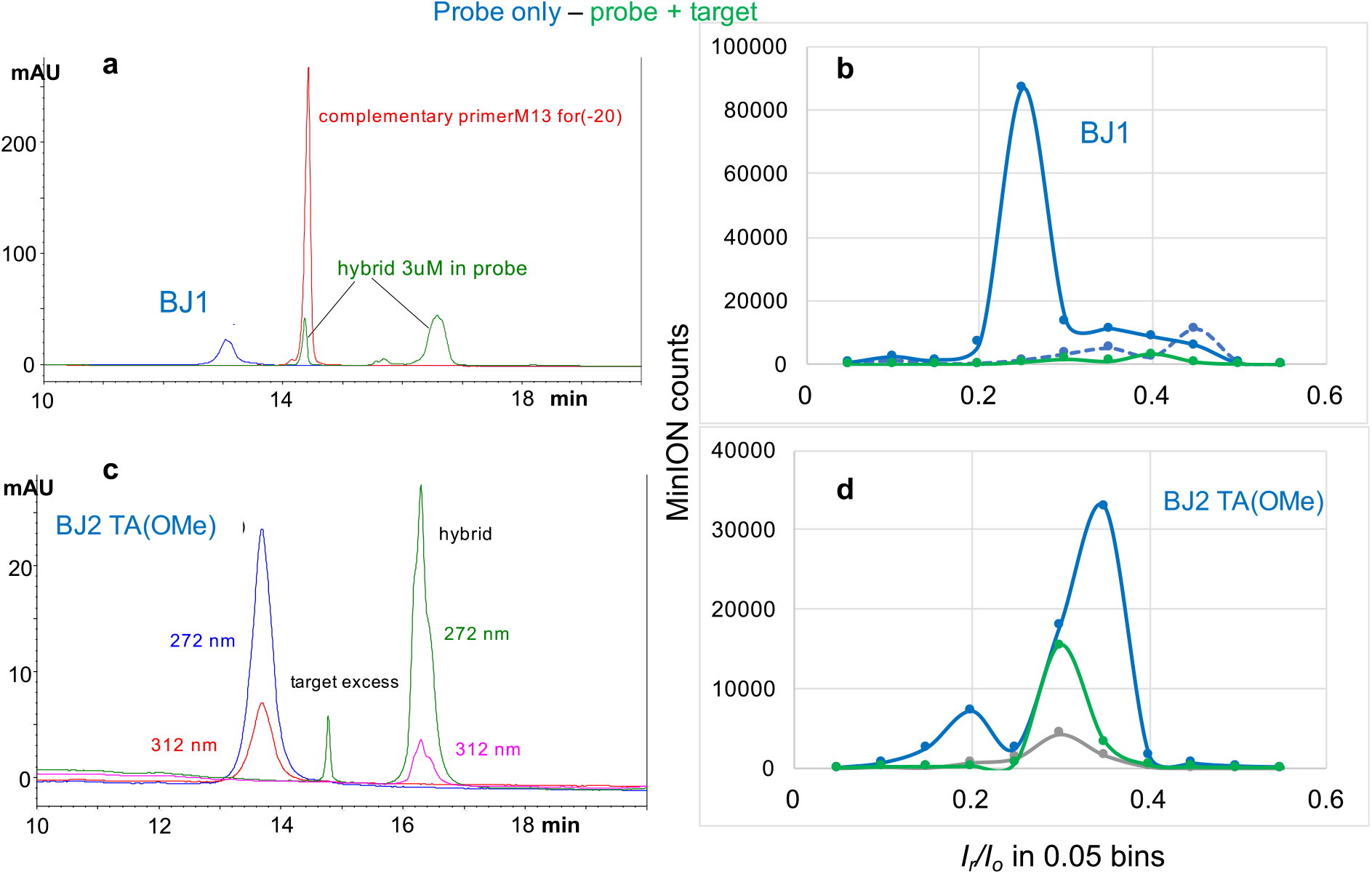
Same samples tested by HPLC and by nanopore; samples in > 90% ONT buffer. **(a)** Overlapping HPLC profiles of samples, at about 0.2nmoles, in ONT buffer (i) probe BJ1 (blue trace), (ii) target (red trace) shown here at a much higher load, and (iii) mixture of probe at about 0.2nmoles and target with about 30% excess over the probe (hybrid, green trace). The HPLC profile of the mixture is consistent with hybridization. **(c)** Overlapping HPLC profiles (i) of the probe BJ2 TA(OMe) at about 0.2nmole load (blue trace) and (ii) its approximately equimolar mixture with the complementary target, complementary primerM13*for*(−41). Target identified as a small peak, at 5% excess over the probe. The HPLC profile of the mixture (hybrid, green trace) is consistent with ds formation (see text). HPLC profiles for these two samples are shown at both the 272 and 312nm (see text and Experimental Section.**(b) and (d)** One hour long nanopore experiments conducted with a MinION flow cell using the same samples as in (a) and (c), respectively; Samples in (b) were used as is, while samples in (d) were used after a 1000-fold or a 100-fold dilution in ONT buffer. **(b)** Probe BJ1 was tested at −180mV (dashed blue trace) and showed few counts. No sample was added, the voltage was raised to −220mV and an additional nanopore experiment was conducted at −220mV (solid blue trace) with counts exceeding 100,000. In contrast to the counts obtained with BJ1, the hybrid sample exhibited very few counts (tested 1h at - 190mV, green trace). **(d)** Nanopore experiments using the same flow cell: (i) control/buffer test (0.75 h tested at −180mV, grey trace), (ii) probe BJ2 TA(OMe) at 0.38pmole load (2h tested at −210mV, blue trace), (iii) equimolar mixture of probe and target at 3.8pmole load each (1h tested at −210mV, green trace). The observation of lower counts with the hybrid sample compared to the probe sample suggest target identification for either (b) and (d), in agreement with the HPLC results. The nanopore experiment with BJ2 TA(OMe) probe at the 0.38pmole load suggests probe detectability at the sub-pmole level. The experiment with the hybrid sample in (d) indicates that the hybrid endures under the experimental conditions of duration and applied voltage (see text). Data acquisition and analysis as described under Fig. 2a.

### Nanopore experiments using the Flongle flow cell

Fig. 2 illustrates results exploiting the Flongle flow cell. Fig. 2a was a first attempt to observe a hybrid. It was known from earlier work with the MinION flow cell that T8, a 32nt RNA with 9 pyrimidines and a total of 8 OsBp tags (see sequence in table), requires high voltage to traverse, exhibits multiple translocation events, and severely obstructs the ionic current exhibiting a maximum of counts at (*I*_*r*_/*I*_*o*_)max≈ 0.1. Repeating the experiment practically duplicated the earlier work on the MinION^49^. Even though not a perfect complement, we used d(TC)_10_ to form the ds complex, as d(CT)_10_ can base-pair 16 out of 20nt, including 8 GC pairs, with T8. While the experiment with probe T8 exhibited, on average, 400 events per channel, the experiment using a 1:1 mixture of probe T8 and d(CT)_10_, yielded less than 50 events per channel for some channels and zero events reported for the rest of the channels (see Figs S1 and S2 in Supplementary Information). Fig. 2b illustrates a test for identification of miRNA122^63^. The probe used here is dmiR122, the exact deoxy complement of miRNA122, and carries 4 T(OsBp) (see table). The sample with probe dmiR122 alone exhibited numerous counts, while the sample with an approximate equimolar mixture of this probe and miRNA122 exhibited markedly fewer counts. A third sample with a 4-times higher miRNA load, composed of miRNA21^64-68^ and miRNA140^63^ also exhibited fewer counts compared to the probe sample. The latter suggests that identification of a target in a complex mixture of miRNAs with excess non-target components is plausible. These as well as other experiments (see later) demonstrate the feasibility of the concept presented in Fig. 1d. They also reveal that both the target and the probe can be either RNA or DNA, the difference being that probes are osmylated oligos, while targets are not. Here we focused on probes that are DNA oligos, because of the lower cost and the higher synthetic product quality compared to RNA oligos.

### Alternative methods to test hybridization between a target and its probe

We sought independent means to test hybridization between osmylated nucleic acids and their DNA or RNA targets. Enzymatic DNA polymerase elongation of an unmodified primer using partially osmylated template ssM13mp18 was our first attempt to obtain support for hybridization, but elongation of primers was not detectable (data not shown). We then tested unmodified ssM13mp18, as the template, and used 30nt long, osmylated at the thymidine (T(OsBp)) primers, BJ1, BJ2, BJ3, and BJ4 (see table). BJ1 carries the identical sequence of primerM13*for*(−20) at the 3’end, and is extended by 13nt at the 5’end. BJ2 carries the identical sequence of primerM13*for*(−41) at the 3’end, and is extended by 6nt at the 5’end. BJ3 and BJ4 have the identical sequence to BJ2 with the exception of one mismatch in the middle of the sequence. Even though BJ2, BJ3, and BJ4 carried 6, 5, and 7 T(OsBp) bases, respectively (table, 3^rd^ column), they were all successfully elongated (Fig. 3a), suggesting that T(OsBp) did not prohibit 1:1 hybridization between intact ssM13mp18 and the probes. In contrast, BJ1, with 6 T(OsBp) did not elongate, presumably due to the presence of a T(OsBp) base at the 3’end and the inability of the enzyme to add a nucleotide to it (Fig. 3a). Still absence of elongation with BJ1 does not necessarily imply absence of hybridization.

These elongation experiments were done at salt concentration presumed to be much lower than the one used for the nanopore experiments, they were limited to ssM13mp18 and to the use of probes with sequences identical to known primers. To extend hybridization tests to miRNAs and any DNA/RNA oligo we developed an HPLC method (see Experimental Section). This HPLC method is based on the HPLC method we use to test for oligo purity with the following modifications: it uses (i) ONT buffer as the sample solvent and (ii) HPLC column temperature at 35°C to mimic the ONT working flow cell temperature. It should be noted that the HPLC column packing may interfere with hybridization, and therefore all the HPLC-based results are purely suggestive. Having said that, there was no instance in our work where the HPLC results and the nanopore experiment contradicted each other. To confirm hybridization HPLC analyses of three separate samples are required. These three samples contain (i) the probe, (ii) the target, and (iii) the sample with the 1:1 mixture of the two components that contains the presumed hybrid (see Figs 3b-d). Absence of hybridization is consistent with an HPLC profile of the mixture sample that closely overlaps with the “sum” of the HPLC profiles of the two components (Fig. 3c). Evidence for hybridization is consistent with a hybrid peak that elutes well resolved from the peaks of the target and of the probe and typically 1 to 1.5 minutes later than either probe or target. In addition, the probe and hybrid peaks exhibit absorbance at 312nm, due to the presence of the OsBp tag, but the target peak does not (see Experimental Section). We intentionally prepared 1:1 mixture samples with a small excess of the target to prevent the probe from being in excess. This is why in many of the HPLC chromatograms the analysis of the hybrid sample includes a smaller peak attributed to the target, in addition to the large peak attributed to the hybrid.

Fig. 3b shows the HPLC chromatograms of samples where both oligos are unmodified nucleic acids. Here the HPLC profile of the sample with the 1:1 mixture is consistent with strong hybridization, based on the features discussed above. Fig. 3d is a repeat of the HPLC analyses in Fig. 3b, only that BJ2 is now osmylated; it is a probe. The corresponding HPLC profile of the 1:1 mixture in Fig. 3d is also consistent with strong hybridization. The same HPLC method indicated that miRNA21 and probe 21EXT (see table) do not hybridize (Fig. 3c). The absence of hybridization in Fig. 3c is attributed to the many OsBp moieties on probe 21EXT, a total of 8 T(OsBp) with 6 T(OsBp) found within the 22nt sequence of the probe which is complementary to the target. Additional hybridization tests confirmed the hypothesis that ds complex formation is prohibited or disfavored in the presence of a large number of OsBp moieties on the probe (see Fig. S3 in the Supplementary Information). Besides the number of OsBp moieties in a molecule, the actual location matters as well. As seen with the BJ1-4 probes, hybridization remains strong, despite a relatively large number of T(OsBp). This is most likely because these tags occupy a small area of the sequence, leaving two rather long subsequences available for duplex formation with the target.

### Hybridization silences the probe in the presence of the DNA oligo target

Experiments in Fig. 2 were conducted using probe, target, and hybrid concentration in the 5μM range. Since sample size of a MinION and that of a Flongle flow cell is 75μL *vs*. 30μL, the 5μM concentration corresponds to a 0.38 *vs*. a 0.15nmole sample load, respectively. It is worth mentioning that sample load for a nanopore experiment is not the same as sample load for an HPLC analysis, as the HPLC injection volume is typically not the same as the flow cell sample size. Fig. 4a illustrates the HPLC hybridization test, described above, using probe BJ1 with 5 T(OsBp) and its target, complementary primer M13*for*(−20). The samples of probe and hybrid were used, as is, for the nanopore experiments, shown in Fig. 4b. We note that probe BJ1 was not enzymatically elongated using ssM13mp18, as the template, which we attributed to the presence of a T(OsBp) base at the 3’end. In Fig. 4 hybridization is documented both by HPLC analysis and by nanopore experiments, as shown by the huge drop in the number of counts reported for the hybrid sample (green trace) compared to the counts reported with the probe sample (solid blue trace). The effect is dramatic for the (*I*_*r*_*/I*_*o*_)_max_, and clearly detectable for the rest of the *I*_*r*_*/I*_*o*_ range.

### Applied Voltage is a critical parameter

Fig. 4b illustrates that testing probe BJ1 with applied voltage at −180mV exhibits very low counts suggesting that probe translocation is inefficient at −180mV (dashed blue trace). Without addition of a new sample, raising the voltage to −210mV (solid blue trace) yielded dramatically more counts compared to the counts obtained at −180m. Fig. S4 in the Supplementary Information illustrates that probes BJ2 and BJ4 follow the same pattern as BJ1. We attribute this observation to the presence of adjacent dT(OsBp) in these probes, and the notion that dT(OsBp) exhibits the lowest observed (*I*_*r*_*/I*_*o*_)_max_, among all tested pyrimidines, a strong indication of heavy crowding. Heavy crowding in adjacent OsBp moieties was concluded from earlier studies using RNAs and the MinION/CsGg^49^, as well as DNAs and the α-Hemolysin nanopores^44^. Since our probes are deoxyoligos and include several T bases in the sequence, the steric hindrance is compounded. Labeled nucleic acids may approach the CsGg pore guided by the applied voltage drop, but to traverse the pore a certain minimum voltage is required. Multiple experiments suggested that the applied voltage for efficient translocation of most of our probes via the proprietary CsGg nanopore is in the range of −210±10mV. In contrast to the observations with the probes, intact DNA oligos exhibit insignificant counts^44,49^ (also data from this study, not shown). Target/unmodified miRNAs exhibit measurable counts with a slight decreasing trend as a function of increasing voltage (Fig. S5 in the Supplemental Information). This is consistent with the expectation that increased voltage results in faster translocation and faster translocation, in turn, results in missed events, as the acquisition rate of the device remains constant at 3 data points per ms. In order to make the counts with native RNAs detectable the total load was 4-times higher than the highest probe load. Experiments with voltage higher than −220mV were not conducted, in order to protect the protein pores that, in our experience, did not last as long at −220mV compared to −180mV. The observation that our probes practically do not traverse the CsGg pore at −180mV is an advantage for a diagnostic test. It provides the opportunity to deplete the sample from excess non-target nucleic acids at −180mV, and then, without adding a new sample, raise the voltage at −220mV in order to detect, or not, the presence of the uncomplexed probe.

### General design for a highly detectable probe

The presence of T(OsBp) moieties in the middle of a sequence is not a feature shared by many potential ctDNA or miRNA targets. Therefore, we designed advanced probes by replacing all dTs in the sequence with dU, modifying some or all the bases as 2’-OMe, added 3 adjacent dTs at the 5’end, and, in some cases added 3 additional dAs at the 3’-end. Addition of dAs at the 3’-end is commonly used to facilitate pore entry^28-30^. Replacing DNA bases with 2’-OMe bases is known to lead to stronger hybridization^69^. Replacing all dTs within the sequence with dUs assures that the presence of OsBp is minimal within the sequence due to the kinetically higher reactivity of T vs. U or dU towards osmylation (see above). This results in the lowest possible number of OsBp moieties within the sequence and the most unhindered hybridization with the target. The addition of 3 adjacent dTs at the 5’end makes the probe undetectable at applied voltage of −180mV and highly detectable at applied voltage of −220mV, as shown by the earlier experiments with the BJ1-4 probes. We then exploited this probe design in nanopore experiments with extra low target loads.

BJ2 TA(OMe) is a probe designed with the above features (see sequence in table). Hybridization between probe BJ2 TA(OMe) and the complementary primerM13*for*(−41) was tested by HPLC at the 5μM concentration range (Fig. 4c). Fig. 4c presents the HPLC analyses of the probe and of the hybrid samples. The target peak in the hybrid sample is easily identified due to its about 10% excess over the probe. The HPLC profiles at both wavelengths, 272nm and 312nm, are shown. Close inspection of these profiles illustrates that the hybrid peak has about half the contribution at 312nm, compared to the corresponding contribution in the probe. This is in agreement with the expectation that the hybrid is half probe and half unmodified target. The actual samples tested by HPLC were diluted with ONT buffer to produce the samples tested by nanopore. The protocol followed for this and all dilutions in this study was done by consecutive 1:10 dilutions with ONT buffer in 0.5mL microcentrifuge tubes. The nanopore experiment with probe BJ2 TA(OMe) was done at a 1000-fold dilution compared to the HPLC sample, i.e. at a 0.38pmole probe load. The corresponding experiment with the hybrid was conducted at 3.8pmole, i.e. at a 10-times higher hybrid load. The higher hybrid load was chosen to test the stability of the hybrid under the experimental parameters especially under the influence of the −220mV applied voltage. Despite the hybrid sample being 10-times more concentrated than the probe, the counts obtained from the hybrid experiment (green trace) are visibly fewer compared to the counts obtained from the probe experiment (blue trace) and suggest that hybrid dissociation is not significant under the tested conditions (Fig. 4d).

### Hybridization silences the probe in the presence of an RNA oligo target

During the development work we experimented with several probe designs. Experiments with two of those designs are illustrated in Fig. 5. Probe 2XdmiR122 is a 44nt oligo with 8T(OsBp) and consists of two fused dmiR122 (see sequence in table). Even though probe dmiR122 exhibited numerous counts at −190mV (Fig. 2b), probe 2XdmiR122 required −220mV (Fig. 5a). The higher voltage is most likely the consequence of heavy crowding within the pore, as 2XdmiR122 incorporates two 4nt groups with 3 OsBp each within a subsequence of 26nt. Efficient hybridization between miRNA122 and 2XdmiR122 was shown by HPLC (Fig. 5c) and confirmed by nanopore, as seen by the huge drop of counts for the hybrid sample (green trace) compared to the probe sample (blue trace, Fig. 5d). The advantage of such fused sequence design lies in exploiting cases where the identical sequence of bases is present in a longer RNA in addition to a miRNA target. Using a probe with a design like the fused 2XdmiR122 may favor hybridization with the miRNA target over the long RNA target, as 2XdmiR122 can form a 44nt ds complex with miRNA122, but only a 22nt ds complex with the longer RNA.

**Fig. 5:**
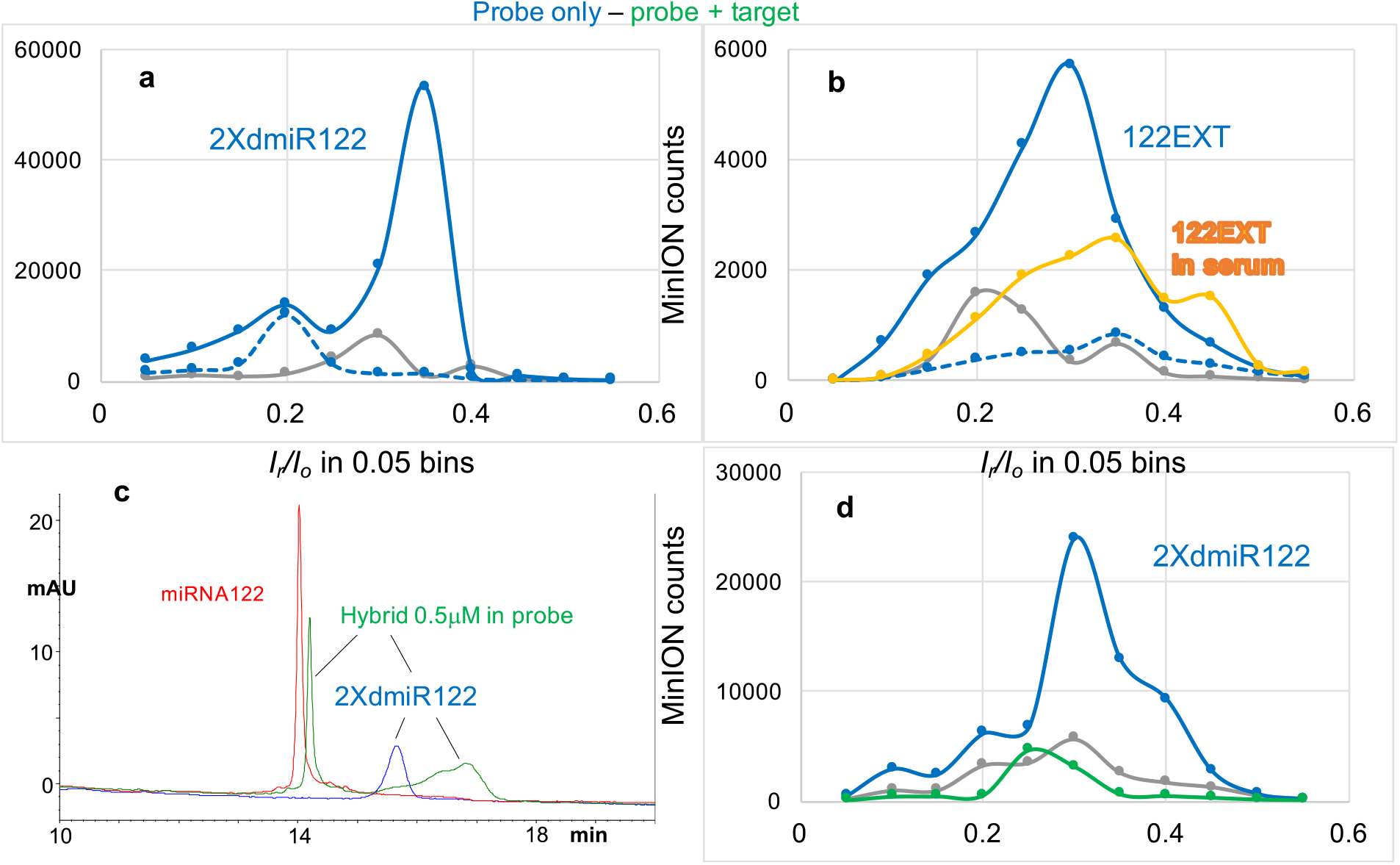
Nanopore experiments testing advanced probe designs and probe detection in 15% human serum - 85% ONT buffer. **(a)** Three consecutive nanopore experiments conducted with the same flow cell. (i) Buffer test 0.75h at −180mV (grey trace), (ii) probe 2XdmiR122 tested 1.5h at −180mV (dashed blue trace) and (iii) no sample added, voltage raised to −220mV and experiment run for 1.5h (solid blue trace). This set of experiments provided solid evidence that, at least, this probe is not traversing the ONT nanopores at −180mV even during a 1.5h long experiment. In addition, it illustrated that this probe traverses the pores under applied voltage of −220mV, and produces a high count of detectable events. **(b)** Four consecutive nanopore experiments conducted with the same but previously used flow cell that carries only around half, i.e. 250, working nanopores. (i) buffer test at −180mV (grey trace), (ii) probe 122EXT at a load 3-times less compared to regular load of 0.38nmoles tested at −180mV (dashed blue trace), (iii) no sample added, voltage raised to −220mV (solid blue trace) and (iv) new sample with same load of probe 122EXT as in (ii), but prepared in 15% human serum - 85% ONT buffer (yellow trace). See discussion in text. **(c)** HPLC profiles of (i) intact miRNA122 (red trace), (ii) probe 2XdmiR122 at 0.5μM, a 10-fold lower concentration compared to our regular 5μM sample concentration (blue trace) and (iii) mixture probe to target=1:2, also at a 0.5μM concentration in probe (hybrid, green trace). All three samples in > 90% ONT buffer and monitored at 260nm using HPLC method B (see Experimental Section). The latter two samples were used, as is, for the nanopore experiments shown in Fig. 5d. **(d)** Consecutive nanopore experiments on the same flow cell. (i) Buffer test, 1h at −220mV (grey trace), (ii) probe 2XdmiR122, 3h at −220mV (blue trace) and (iii) hybrid sample with miRNA122, in 2-fold excess over the probe, tested for 1h at −220mV (green trace). This experiment confirms sensitivity in the detection of 2XdmiR122, albeit via a 3h long experiment, and confirms that hybridization with target results in severely reduced counts, i.e. silencing. Data acquisition and analysis for the nanopore experiments as described under Fig. 2a.

Fig. 5b presents an additional probe design, exemplified by probe 122EXT (sequence in table). Probe 122EXT has the identical sequence of dmiR122 with the addition of 3 adjacent dTs at the 5’-end. This probe requires −220mV to show numerous counts (blue solid trace), as seen by comparison to the nanopore experiment at −180mV (dashed blue trace). A nanopore experiment conducted with probe 122EXT in a sample prepared in 15% human serum and 85% ONT buffer (yellow trace) exhibits about half the counts compared to the sample prepared in over 95% ONT buffer (blue solid trace). The reduction in counts may be attributed to the lower ionic strength due to the presence of serum, and/or to the aged flow cell and/or to serum interference. Despite the lower counts probe 122EXT is easily discriminated from the control/buffer test (grey trace) indicating the unhindered probe detection in an unknown sample that contains a body fluid, such as human serum.

miRNA21 is an important biomarker for a number of diseases^64-68^, so its identification was our priority. HPLC tests with miRNA21 and probes dmiR21 (not shown) or 21EXT indicated no detectable hybridization with miRNA21 (Fig. S6 in the Supplementary Information). Inability to form the hybrid was attributed to the presence of more than 5 T(OsBp) moieties and the fact that these moieties are spread over the 22nt sequence. Advanced probe design led to probes that efficiently hybridized with miRNA21. Probe dmiR21(OMe) is a 22nt oligo, complementary to miRNA21, where all bases are 2’-OMe, dTs are replaced with mU, and osmylation resulted in the addition of 2.85 OsBp moieties, on average, per molecule (see table and Experimental Section). Please note that mU is 2’-OMe-U and 2’-OMe-base-deoxyribose is the same chemical as 2’-OMe-base-ribose. The osmylated product is a mixture containing primarily molecules with 2 or 3 OsBp moieties, and also molecules containing OsBp moieties at different bases (called here topoisomers)^49,58^. Our chromatography resolves molecules that carry one, two or three OsBp moieties and often resolves topoisomers too^59^. This is why the HPLC profile of the probe consists of two separate peaks attributed to a molecule with 2 tags and to a molecule with 3 tags (blue trace, Fig. 6a). Similarly, the HPLC profile of the hybrid appears as multiple peaks (red trace, Fig. 6a). We tested by HPLC the stability of RNAs (see below) and the stability of the hybrid of probe dmiR21(OMe) with miRNA21 in a solvent composed of 15% human serum and 85% ONT buffer. Fig. S7b illustrates that the tested RNAs, i.e. miRNA140 and the 100nt RNA, both degraded within minutes, whereas the hybrid peark remained practically unchanged, suggesting that the hybrids formed between our probes and their RNA targets are expected to be stable during the duration of a nanopore experiment in human serum/ONT buffer.

**Fig. 6:**
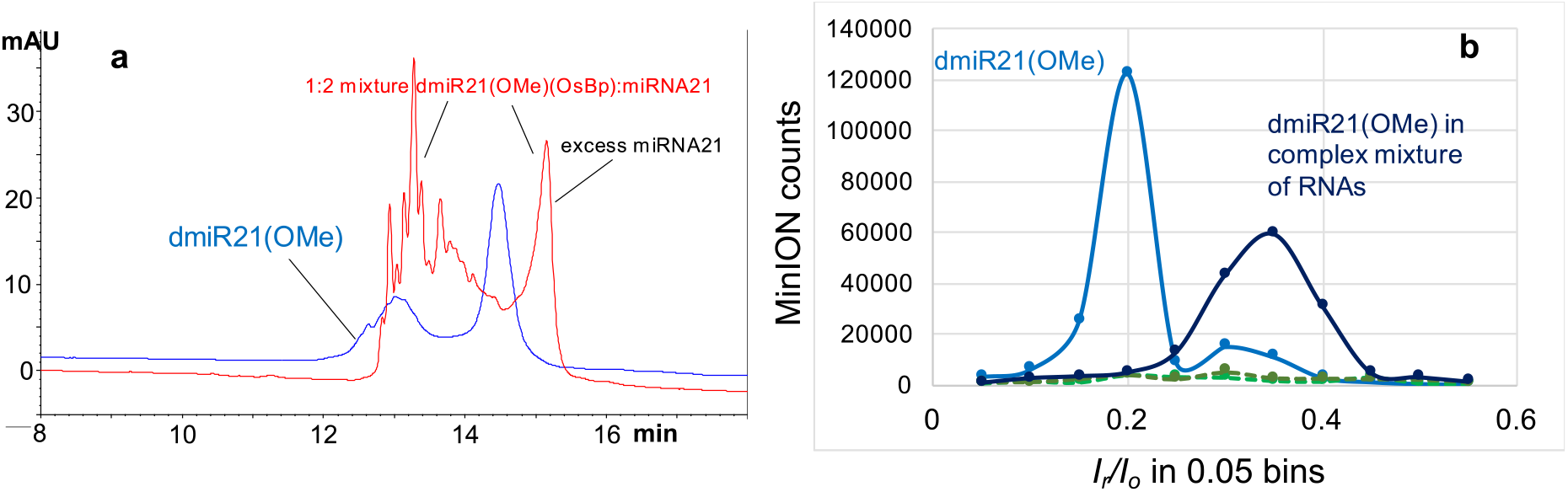
Targeting miRNA21 in a complex mixture. **(a)** HPLC profiles of two samples analyzed with HPLC method B (see Experimental Section): (i) probe dmiR21(OMe) at 0.15nmole load and (ii) 1:2 mixture of this probe with miRNA21 at a 0.30nmole probe load. The probe’s HPLC profile exhibits two peaks, the larger one eluting after the minor one (blue trace). This profile is consistent with the determined value of 2.85 OsBp moieties per molecule, on average, meaning that this preparation includes molecules with 2 and molecules with 3 OsBp tags. The HPLC profile of the 1:2 mixture of probe to target exhibits a single rather sharp peak eluting after a broad rather complex peak (red trace). We confirmed that the sharp peak corresponds to the excess miRNA21 target and attribute the broad complex peak to multiple hybrids, the result of one target and many probes, all complementary to the target, but each one of them carrying OsBp moieties at a different nucleobase. It is reasonable to assume that the chromatography resolves these hybrids, as it resolves topoisomers with such short osmylated oligos^59^ (HPLC method B in Experimental Section). The observation of distinct HPLC profiles between probe and mixture samples is consistent with hybridization. **(b)** Four nanopore MinION experiments two of them using the exact samples analyzed by HPLC in (a): (i) probe dmiR21(OMe), tested for 2h at −180mV, exhibited over 100,000 events (light blue solid trace), (ii) 1:2 mixture of this probe with miRNA21, tested for 1h at −180mV, exhibited negligible counts (hybrid, light green dashed trace). Two additional experiments tested the effect of excess non-target RNA in the translocation properties of probe dmiR21(OMe) and its hybrid with miRNA21. The excess non-target RNA was at a 10-fold higher load compared to the probe and was composed of equimolar amount of miRNA140 and a 100nt long RNA. Nanopore experiments in the presence of excess non-target RNA (iii) hybrid with miRNA21, 1h at - 180mV (dark green dashed trace), and (iv) probe dmiR21(OMe), 2h at −200mV (dark blue solid trace). Higher voltage used here to compensate for the hours that this flow cell had worked already, per ONT protocol. The effect of the excess non-target RNA on the hybrid, if any, is not detectable, as the counts are too few. The effect of the excess non-target RNA on the probe appears to be a profile shift and reduced counts by a factor of 2, even though this reduction can be attributed to the number of working nanopores that is also reduced by a factor of about 2. Please note that probe dmiR21(OMe) translocates efficiently at an applied voltage of −180mV, as it does not contain any adjacent osmylated dTs. Data acquisition and analysis as described under Fig. 2a.

Hybridization is consistent with the distinct HPLC profiles observed with the probe and the hybrid samples (Fig. 6a). Because dmiR21(OMe) does not contain 3 adjacent T(OsBp), applied voltage at - 180mV was sufficient to thread this probe via the pore. Plenty of events were reported with a 0.75nmole probe sample and markedly fewer with a 1.5nmole hybrid sample (Fig. 6b, compare solid light blue trace with dashed light blue trace). Counts with the hybrid appear comparable to counts obtained with buffer (not shown). Two additional nanopore experiments were conducted with the same probe and hybrid loads, as before, but in the presence of excess non-target RNA components. These components were miRNA140, and a 100nt RNA, at a total 10-fold higher load compared to the probe. The nanopore profiles of the probe samples with or without the excess non-target RNA are distinct suggesting influence by the excess material and/or an aged flow cell (light blue vs. dark blue solid traces). Still the two experiments in the presence of the non-target RNA show efficient identification of the target by comparing the numerous counts of the probe sample (dark blue solid trace) to the few counts of the hybrid sample (dark blue dashed trace). This discrimination suggests that the presence of non-target miRNAs and longer RNAs in a complex mixture do not prevent target identification.

### Due to the importance of miRNAs as biomarkers^63-68^

we conducted experiments at extra low load with a probe design that is broadly applicable to any target sequence and exhibits optimal translocation properties. We selected probes 140EXTmU and 21EXTmU (see sequences in table) to target miRNA140 and miRNA21, respectively. These probes have a sequence complementary to their target, mU replacing T within the sequence, 3 additional adjacent dTs at the 5’-end, and 2 or 3 additional dAs at the 3’end. These oligos were osmylated using the validated labeling process that adds, on average, 4 to 5 OsBp tags per molecule, 3 of which occupy the 5’-end, and the other 1 or 2 are randomly allocated within the sequence (see table and Experimental Section). Due to the heavy crowding at the one end of the probe, applied voltage in the range of −210±10mV is required for probe efficient translocation and detection (see discussion above). Fig. 7a illustrates detection with excellent sensitivity at the 47fmole level with 140EXT(mU), and Fig. 7b illustrates detection of this probe at an even higher sensitivity of 3.5amole probe load. Identification of the target miRNA140 is also evident by comparing the probe (blue trace) to the hybrid (green trace) with hybrid load also at 3.5amole. Fig. 7c shows the HPLC profile of the hybrid sample between probe 21EXT(mU) and miRNA21, which was then diluted by a 3×10^8^ factor for the nanopore experiment to test the hybrid at the 2.7amole level (green trace in Fig. 7d). The 21EXT(mU) probe sample (HPLC profile not shown) was diluted by a 1×10^9^ factor in order to conduct the nanopore experiment at the 0.9amole level of (blue trace in Fig. 7d) which is the lowest load tested in this study. Identification of miRNA21 is evident in Fig. 7b by visual comparison of the probe counts (blue trace) to the counts of the hybrid (green trace). The reason that the counts observed with the 140EXT(mU) probe (top reaching 6,000) are fewer compared to the counts observed with the 21EXT(mU) probe (top reaching 24,000) is, at least partially, due to an aged flow cell that had only about 30% of working pores. Whether or not proportionality in probe counts can be obtained from the ONT/OsBp platform for a specific probe was not tested here, primarily because probe concentration is a known quantity. Most importantly, identification of the target is based on a nanopore profile that exhibits an insignificant number of counts, also comparable to the counts of the control/buffer experiment. The main reason we tested every probe by nanopore, is because we wanted to compare different probe designs and confirm high detectability and high sensitivity.

**Fig. 7:**
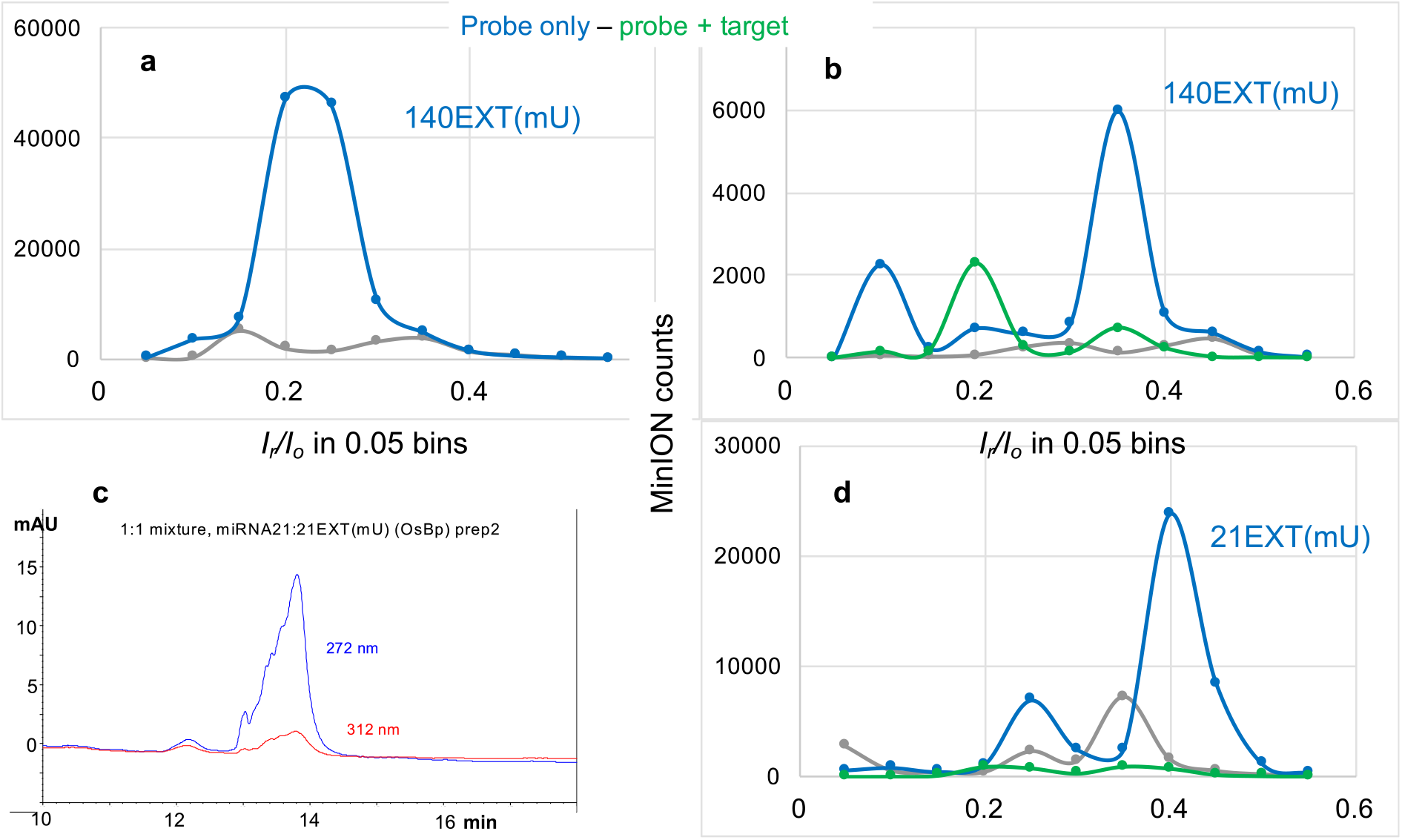
Nanopore experiments targeting miRNA140 or miRNA21 at single-digit attomole load. **(a)** Consecutive nanopore experiments conducted with the same flow cell: (i) buffer test, 1h at −210mV (grey trace), (ii) probe 140EXT(mU) at a 47fmole load, 1.5h at −210 mV. **(b)** Consecutive nanopore experiments conducted with the same used flow cell with less than 200 working nanopores: (i) buffer test, 1h at −210mV (grey trace), (ii) probe 140EXT(mU) at a 3.5amole load, 1.5h at −210 mV (blue trace) and (iii) equimolar mixture of 140EXT(mU) and miRNA140, at 3.5amole load each, 1.5h at −210 mV (hybrid, green trace). **(c)** HPLC profile of a sample with equimolar concentration of probe 21EXT(mU) and target miRNA21 using HPLC method B (Experimental Section). The appearance of a single peak is consistent with hybridization. Sample monitored at two different wavelengths to show the absorbance at 312nm attributed to the presence of the osmylated probe. **(d)** Consecutive nanopore experiments with the same flow cell; samples were first tested by HPLC, then diluted with ONT buffer by a factor of 10^9^ or 3×10^8^ for probe or hybrid, respectively (see text and Experimental Section). (i) buffer test, (grey trace), (ii) probe 21EX(mU) at 0.9amole, (blue trace), and (iii) 1:1 mixture of probe 21EXT(mU) with miRNA21 at 2.8amole load each, (hybrid, green trace). Hybrid exhibits dramatically fewer counts compared to the probe. Data acquisition and analysis for the nanopore experiments as described under Fig. 2a.

### The potential of the ONT/OsBp nanopore platform

The above experiments range in probe load from 0.38nmole to 0.9amole, practically <u>spanning 9 orders of magnitude</u>. In this range evidence is presented for probe detection, and clear distinction between the sample that contains the target and the sample that doesn’t. We propose as lower detectability limit, 3-times higher count of events with probe alone compared to the counts of the hybrid. Proportionality between the count of events and the probe concentration was not tested here, as the duration of the experiments remains in the range of 1 to 3h and does not reflect the sample load variation. Since our test does not detect the hybrid and therefore does not measure the hybrid molecules, quantification of the target depends on the known probe load. The test requires an estimate for target load and depending on the outcome of the first experiment, one can vary the probe load by a factor of 5 below or above in the next experiment. We estimate that target quantification can be attained with about 30% accuracy. Here we explored a number of probe designs and proposed one design that can practically match any ssDNA or ssRNA oligo target. This type of probe, with examples identified as 140EXT(mU) and 21EXT(mU), exhibited high sensitivity at the amole load level. No attempt was made to test for specificity. Discrimination between one target and another target with similar sequence will need to be addressed case-by-case. The HPLC method developed here can evaluate hybridization between a target and a tentative probe and it can also assess probe hybridization extent and tentative discrimination between two targets with similar sequence. Here the flow cell temperature was at the factory preset temperature, but nanopore proteins are known to be stable in a certain temperature range. Flow cell temperature, if left at the discretion of the user, can provide a means to improve specificity. We also did not replace the proprietary ONT Flush buffer. The latter was developed for sequencing, and a different buffer may be more suitable for our application.

Preliminary experiments indicate that the hybrid of miRNA21 with dmiR21(OMe) is relatively stable in a 15%-85% serum-ONT buffer and that probe 122EXT is detectable by nanopore in a 15%-85% serum-ONT buffer. These experiments suggest that using the ONT/OsBp nanopore platform with blood serum samples is feasible. Considering that the MinION flow cell uses 75μL sample volume and 15% could be blood serum, then approximately 11μL of a human serum sample can be directly tested in a nanopore experiment. Assuming that an 11μL human serum sample contains about 3amole of miRNA21, miRNA140, or tentatively any other miRNA, our general probe design should be able to detect it, and as a corollary detect the presence/absence of the target. In this context, our test qualifies as a follow-up, point-of-care diagnostic test.

Even though not tested here, RNA-based probes may successfully target a relatively short dsDNA after addition of the probe to the unknown sample and denaturation. Such identification may be feasible, as RNA/DNA hybrids are known to be more stable compared to DNA/DNA hybrids. RNA probes exhibit a different nanopore profile compared to DNA-based probes, as indicated by comparison of the (*I*_*r*_*/I*_*o*_)_max_ in Fig. 2a with the corresponding (*I*_*r*_*/I*_*o*_)_max_ in all the other figures representing probes. This feature can lead to probe multiplexing in order to test more than one target at a time. Similarly, probes like dmiR21(OMe) that are missing the 3 adjacent dTs at the 5’-end were seen to translocate at −180mV, while probes with 3 adjacent dTs require −220mV. This distinction may be also exploited in order to multiplex probes. Distinct nanopore profiles will reveal which one of the probes is silenced. With just one probe, the test can conclusively identify the presence/absence of the target by comparing the total count of events of the probe alone to the total count of events to the mixture of the probe with the unknown sample. With a multiplexed test, counts should be plotted as histograms, just as presented here, in order to determine which probe is missing.

## CONCLUSIONS

We presented ion-channel single molecule experiments conducted using portable, commercially available nanopore devices from ONT. While a number of experimental nanopore platforms paved the way for using nanopores as single molecule sensors, our work appears to be the first to explore ONT devices. The proposed test targets DNA and RNA oligos and exhibits a 9 orders of magnitude range of detection, approaching single-digit attomole target sensitivity from an 11μL sample. Our technology exploits the relatively slow data acquisition rate of the ONT platform in which native oligos traverse through the nanopores too rapidly to be detected, while probes selectively tagged with a bulky Osmium label (OsBp) traverse at a slower, detectable rate. Furthermore, OsBp modified probes hybridized to a target oligo do not enter the pore, effectively “silencing” the probe by quantitatively reducing the number of detected events. This silencing allows quantification of the target based on the known concentration of the probe. These properties enable detection and quantification of highly dilute samples such as ctDNA and miRNAs found in bodily fluids.

Expense and product quality considerations led us to select DNA over RNA oligos as probes. Our experiments revolved around optimizing the probe’s design to make it of general applicability, and aptly detectable by the proprietary ONT/CsGg nanopore. A probe that successfully identified miRNA targets at the single-digit attomole level has sequence complementary to the target, 2’-OMe-U replacing dT within the sequence, 3 additional adjacent dT residues at the 5’-end, and 3 added dA residues at the 3’-end. Additionally, the oligo is osmylated to add 4 to 5 OsBp tags per molecule, 3 of which occupy the 5’-end, and the other 1 or 2 are randomly allocated within the sequence. Due to the heavy crowding at the one end of the probe, applied voltage in the range of −210±10mV is required for probe translocation and detection. This feature is advantageous, as it allows for depletion of non-target material at −180mV prior to performing the diagnostic experiment at −210mV. Probes may also be multiplexed and target up to 3 oligos when using probe designs that exhibit distinct nanopore profiles. Preliminary experiments in 15% human serum suggest that probes and the resulting hybrids are stable in such medium, and show the feasibility for identifying short DNA and RNA from body fluid samples using the ONT/OsBp nanopore platform.

## EXPERIMENTAL SECTION

### RNAs and other Reagents

Custom-made RNA oligos were purchased from Dharmacon (Horizon Discovery Group). Custom-made deoxyoligos were purchased from Integrated DNA Technologies (IDT). Sequences and UV/Vis properties of their osmylated derivatives are listed in the table. Purity of oligos was tested by HPLC in-house and typically found to be >85%. Oligos were diluted with Ambion Nuclease-free water, not DEPC treated, from Thermo Fisher Scientific typically to 100 or 200 μM stock solutions and stored at −20°C. HPLC profiles from the analyses of both the intact and the osmylated oligo are included in the Supplementary Section. Buffer DNase-free and RNAse-free TRIS.HCl 1.0 M pH 8.0 Ultrapure was purchased from Invitrogen and used to prepare the HPLC mobile phase. NaCl crystalline ACS min 99.0% from Alfa Aesar. Distilled water from Alhambra was used for preparation of HPLC mobile phase. A 4% aqueous osmium tetroxide solution (0.1575 M OsO_4_ in ampules at 2 mL each) was purchased from Electron Microscopy Sciences. 2,2’-Bipyridine 99+% (bipy) was purchased from Acros Organics. Human Serum from human male AB plasma and NaOH 1N Bioreagent were purchased from Sigma. ss M13mp18, primer M13(fwd6097), NEBuffer 2.1, and Klenow Fragment of DNA Polymerase I (3’-5 exo)(M0212) were kindly provided by New England Biolabs, Ipswich, MA, USA.

*(Pyrimidine)OsBp is a chromophore*. Selective labeling of a nucleic acid requires an assay for quality control. Addition of OsBp to the C5-C6 Py double bond and formation of Py(OsBp) creates a new chromophore in the wavelength range of 300 to 320nm^54^, where nucleic acids exhibit negligible absorbance. We exploited this observation and used a deoxyoligo training set to show that extent of osmylation can be measured using the equation: R(312/272) = 2x(No of osmylated pyrimidines/total nt of nucleotides)^54^. Value R(312/272) is the ratio of the observed peak absorbance at 312nm over the observed peak absorbance at 272nm (the peak shape could be sharp or broad or multiple peaks)^54,58,59^. The wavelengths 312 nm and 272 nm were chosen in order to maximize the effect and to equalize contributions by different pyrimidines^54^. Absorbance at 272nm is about 75% of the absorbance at 260nm practically for either intact or osmylated nucleic acids. Using the ratio R instead of the absorbance at 312nm normalizes the measurement, and minimizes instrument sampling variation.

As deduced by experimenting with an oligo training set, when observed value R(312/272) = 2x(# of pyrimidines/total # of nucleotides), osmylation is practically 100% complete and all the pyrimidines carry one OsBp moiety ^54,58^. When observed R(312/272) < 2x(# of pyrimidines/total # of nucleotides), osmylation is partial and the number of osmylated pyrimidines or OsBp moieties can be calculated based on the same equation (see columns # 5 and 7 in the table). With partial osmylation the number obtained from the equation refers to OsBp moieties, on average. Molecules carry an integer number of OsBp moieties and therefore some molecules will have fewer, and some molecules will have more than the calculated value distributed in a statistically unbiased manner. We have shown that osmylation occurs randomly, but depends on the relative reactivity of OsBp for the pyrimidines. Relative reactivity towards osmylation, determined by kinetic measurements at 26°C in water using deoxyribooligos^44^ is T/dC=28, dU/dC=3.75 and therefore T/dU=7.5, in agreement with earlier studies^58^. Comparable kinetic measurements of OsBp reactivity towards ribooligos determined U/C=4.7 and 5-MeU/C=44^49^; hence 5-MeU/U=9.3, with 5-MeU carrying the identical nucleobase as T. These measurements suggest that reactivity of OsBp towards a nucleotide is primarily determined by the identity of the nucleobase, while the sugar and the backbone make an insignificant contribution. Because of the markedly higher reactivity of OsBp towards T, compared to U and C, conditions can be found to osmylate practically 100% all dTs, while some dU and very few dC become osmylated^54,58^. We have exploited this dramatically higher reactivity, added 3 dTs at the 5’end of a probe, replaced the dTs in the sequence with dU or mU, in order to keep the base-pairing with Adenosine in the target intact, and optimized the manufacturing process as detailed below.

### Manufacturing of OsBp-nucleic acids

OsBp reagent is prepared by weighing the equivalent of 15.7mM of bipy (49.2mg) in a 20mL scintillation vial, adding 18μL of water and stirring at room temperature until bipy dissolves, followed by transferring the full content of a 2μL 4% OsO_4_ solution supplied in an ampule. Dissolving bipy in water before the addition of OsO_4_ (protocols a, b, c, and d) results in a more consistent and potent preparation compared to dissolving bipy after the addition of OsO_4_ (protocol o). The transfer should be done using a glass pipette inside a safety hood^70^. The resulting solution is an aqueous 20mL 15.75 mM OsBp (0.4%) stock solution, equimolar in OsO_4_ and bipy. The concentration of the OsBp stock solution is limited by the solubility of bipy in water and adding OsO_4_ does not increase it, as the complex has a low association constant. OsBp complex represents an approximate 5% of the total, as measured by CE^58^. Care should be taken that this preparation, as well as any other work using OsBp is conducted in stoppered glass vials in a well-ventilated area. Leftover solutions of OsO_4_ and/or OsBp may be mixed with corn oil to neutralize unreacted OsO_4_ and properly discarded based on specific local regulations^70^. The freshly prepared OsBp stock solution is dispensed in HPLC vials and kept at - 20°C. Each vial can be stored at 4°C and used for a month without loss of potency; typical pipette tips can be used for manufacturing of OsBp-labeled nucleic acids. OsBp stock solutions should be validated before first use by running a known reaction. For osmylation reactions a 20-fold excess of OsBp over the reactive pyrimidine in monomer equivalents is required to ensure pseudo-first order kinetics, and to assure successful use of protocols. Manufacturing conditions, i.e. OsBp concentration and labeling duration varies significantly depending on the presence of T, and the desired result of osmylating all pyrimidines, T only, or just a fraction of C and U in RNA. For the purpose of this study, when T-osmylation is required protocol b is recommended and when partial osmylation of U and C is required protocol c is recommended. These choices were facilitated by testing additional protocols (o and a), as identified in the table. Protocol d is used when the probe carries no Ts. Quenching of the osmylation reaction occurs upon purification. Purification from excess OsBp was done with spin columns (TC-100 FC from TrimGen Corporation) according to the manufacturer’s instructions. Briefly, spin columns are filled with the manufacturer’s proprietary solution and centrifuged at 4,000 rpm for 4 min; the resulting solution and the microcentrifuge tube are discarded. Then 40 to 120μL of an osmylation reaction mixture is transferred to the spin column and centrifuged at 4,000 rpm for 4 min using a clean microcentrifuge tube. The centrifuged solution is the purified osmylated oligo. This purification method retains the volume/concentration of the sample, and close to 100% recovery of RNA is achieved. For purity and characterization of the intact and osmylated oligos see figures S8-S18 in the Supplementary Information.

Recommended protocol b for T-osmylation is 30±2min incubation with 2.6mM OsBp in water at room temperature (see table under ^b^). Recommended protocol c for partial U- and C-osmylation of probes that do not carry dTs is 30±2min with 3.9mM OsBp in water at room temperature (see table under ^c^). There were two additional protocols tested, but found to be less optimal: Protocol where bipy was not dissolved in water prior to the OsO_4_ addition and used 40 min incubation with 2.6mM OsBp (see table under °) and the protocol using 45 min incubation with 2.6mM OsBp (see table under ^a^). The presence of 2’-OMe groups does not affect markedly the extent of osmylation, as seen by comparable osmylation extent for 100nt RNA and 100nt RNA(2’-OMe) that carries about 50% 2’-OMe bases. What affects osmylation extent is the number of U in the oligo as seen in Table S1 and Fig. S19 in the Supplementary Information. This is because the reactivity of OsBp towards U is 4.7 times higher compared to the reactivity towards C. Because the osmylation protocols do not go to completion U(OsBp) or mU(OsBp) are kinetically preferred over C(OsBp). With protocol^b^ osmylation of other pyrimidines is negligible compared to T-osmylation, but with protocol^c^ the extent of osmylation towards U and C is measurable (Table S1 in the Supplementary Information). Fig. S19 suggests a linear correlation between the number of OsBp moieties and the number of U present in an oligo spanning from 22nt to 100nt oligos. One may use the slope=0.43 of the graph to estimate extent of osmylation with protocol^c^ for any oligo based on the number of Us in the sequence.

We used 2mL HPLC vials fitted with 120μL glass inserts to carry the manufacturing reactions. Removing the reaction product from these inserts and transferring it onto the purification spin column requires a long and narrow 20μL pipette tip. Osmylated nucleic acids are as stable as the corresponding nucleic acid, and the OsBp label is unreactive. They can be stored in the 1.5mL microcentrifuge tubes, at −20°C for years. The concentrations of 2.6, 3.9 and 5.2mM correspond to 1/6, 1/4 and 1/3 dilutions of the 15.75 mM OsBp stock solution, respectively. Deviations from these two protocols are included in the table, identified as protocols ° and ^a^. An additional protocol (^d^) was used in order to achieve higher extent of osmylation with probes that do not contain any T, such as dmiR21(OMe) and make it a detectable probe.

### Enzymatic elongation reactions

The ability of DNA polymerase to extend an osmylated primer was examined *in vitro* using ssM13mp18 annealed to unmodified and osmylated oligonucleotides. ssM13mp18 at a concentration of 42 nM was mixed with 0.42 µM primer in NEBuffer 2.1(New England Biolabs). Samples were heated to 90°C for 30 seconds and cooled to 25°C at 0.1°C/sec. Polymerization reactions contained these annealed complexes: 8.4 nM M13mp18, 1.25X NEBuffer 2.1 (NEBuffer is 10 mM TrisHCl pH 7.5, 50mM NaCl, 1mM EDTA), 0.25 mM each of dGTP, dATP, and dTTP, 0.025 mM α-[^32^P] dCTP, and 7U/mL Klenow Fragment of DNA Polymerase I (3’-5 exo-) (NEB, M0212). Reactions were incubated at 37°C and incorporation of labeled dCMP was monitored by an acid precipitation assay. Time points were taken at 5, 10, and 20 minutes.

As seen in Fig. 3a, no incorporation was noted when no oligonucleotide was added, or if the oligonucleotide, primerM13*rev*(−48), was not complementary to the ssM13mp18. In contrast, most oligonucleotides predicted to anneal to the DNA template gave robust incorporation, even when osmylated, with maximal incorporation corresponding to roughly one round of replication on the M13mp18 DNA template. Incorporation noted for M13(fwd6097), BJ2, BJ3, and BJ4 were equivalent despite interior single base mismatches within BJ3 and BJ4 (see sequences in table). Despite overall equivalent osmylation levels for BJ2, BJ3 and BJ4, markedly lower levels of incorporation were noted for BJ1, most likely due to presence of OsBp at the terminal 3’-T(OsBp) residue. Control experiments mixing BJ1 with M13(fwd6097) displayed full incorporation, discounting soluble inhibitors as the cause of low incorporation with BJ1 (data not shown).

### HPLC methods

The HPLC method to assess oligo purity was developed earlier, and used here to assess purity of the oligos listed in the table. This method was optimized (see below) and used to assess hybridization between two oligos in a mixture; validation of this method was conducted using a 1:1 mixture of intact oligos known to hybridize (see Fig. 3b). Analyses were conducted automatically using a thermostatted autosampler. HPLC peaks were detected and identified using a diode array detector (DAD) in the UV–vis region 200–450 nm. The chromatograms were recorded at 260, 272, and 312nm and reported here selectively. Samples were prepared with RNAse free water, but buffers were not. For HPLC analysis we used an Agilent 1100/1200 LC HPLC equipped with a binary pump, Diode Array Detector (DAD), a 1290 Infinity Autosampler/Thermostat, and Chemstation software Rev.B.04.01 SP1 for data acquisition and processing. For sample analyses we used IEX HPLC column DNAPac PA200 from ThermoFisher Scientific (Dionex) in a 2×250mm configuration. The performance of the instrument and the column was qualified using standards every time ahead as well as after analysis of research samples. The HPLC method was developed to assess hybridization in a sample approximately 90% in aqueous ONT buffer at pH 8 and column thermostat at 35°C. This HPLC method (identified as B) uses the DNAPac PA200 column with a 0.45mL/min flow, mobile phase A (MPA) aqueous 25mM TRIS.HCl pH 8 buffer, mobile phase B (MPB) aqueous 1.5M NaCl in a 25mM TRIS.HCl pH 8 buffer, a gradient from 10% MPB to 50% MPB in 20min and an additional 10min for wash and equilibration to initial conditions, i.e. 90% MPA. Column temperature was set at 35°C to mimic the flow cell temperature. To include the 100nt RNA the chromatography was modified to HPLC Method C. Specifically the gradient was made steeper, 10% MPB to 75% MPB in 20min, and everything else remained as is. Because of the approximately neutral pH the long 100nt RNA elutes as a broad peak resembling a mixture. This is because aqueous pH 8 does not denature the various conformations of a long RNA, as reported earlier^61^. ONT buffer is proprietary material, and its composition is not known to us. ONT buffer has a UV-Vis component that in this chromatography elutes in the void volume and does not interfere with the analysis of the samples. Sample injection volume was typically at 5μL, and not higher than 10μL. Please note that for the hybridization test to work this chromatography should be used in conjunction with samples in ONT buffer or any other medium that favors complexation. Some of the oligos and their osmylated derivatives were analyzed using a method idenfied as HPLC method A, which is recommended for oligo purity analysis^61^, but not suitable to test hybridization. HPLC method A uses the DNAPac PA200 column with a 0.45mL/min flow, and column temperature at 30°C. Mobile phase A (MPA) aqueous pH 12.0±0.2 with 0.01 N NaOH, mobile phase B (MPB) aqueous 1.5M NaCl in pH 12.0±0.2 with 0.01 N NaOH, a gradient from 20% MPB to 95% MPB in 12min and an additional 8min for wash and equilibration to initial conditions, i.e. 80% MPA.

### Single molecule ion-channel conductance experiments with the CsGg nanopore in a MinION or a Flongle (ONT platform)

ONT instructions were followed, as needed. Briefly, a new flow cell is flushed with ONT Flush buffer via the priming port in order to remove the storage solution. The sample is added via the sample port, while the priming port is open. Air bubbles from the flow cell are removed from the priming port right before every use. We stored the flow cells in ONT Flush buffer for up to 2 months, while using them as needed. The MinION takes a 75μL sample and the Flongle a 30μL sample. Flongle flow cells require an adaptor but work on the same device as the MinION flow cells. The software MINKNOW to run the nanopore experiments was downloaded on a MacBook Pro laptop used for these experiments. All the functions necessary to test the flow cell and run the experiments are done via the MINKNOW software tool. Raw data files were acquired in *fast-5* format, which were then analyzed by the *OsBp_detect* software. Size of fast-5 files for the experiments depend on the flow cell and experiment duration and vary between 1.5 and 6GB. *Fast-5* files can be directly visualized in MatLab (from Mathworks) 2D format, once the experiment is completed. MINKNOW allows for monitoring in real time any chosen channel, up to 10 channels, so one doesn’t need to wait for the experiment to be done to see the *i-t* recordings.

### Samples were either intact oligos or Yenos proprietary osmylated RNA oligos, or mixtures thereof

Concentrations were typically at or below 10 μM oligo in no less than 80% of ONT buffer. No library was prepared, and no processing enzyme was added, such that all the translocations reported here are unassisted and voltage-driven. Experiments lasted no longer than 1.5h, but the same experiment could be extended by stopping it, and restarting it later, or next day without adding a new sample. Running more than 2 experiments per day on the same flow cell was avoided. In most cases the first experiment was a “buffer or control test” to assess the flow cell’s baseline. We were surprised on how many events are produced by the ONT buffer and anticipate that ONT may be able to recommend a more suitable buffer for our applications. Duration of the buffer test was kept as short as possible, because the nanopores do not last for more than 15 hours under our experimental conditions. Per ONT protocol, applied voltage was raised by about 10mV for every 5 working hours, in an attempt to keep the open pore ionic current (*I*_*o*_) constant. This is why experiments compared to one another are done at seemingly different applied voltage. The MinION flow cell has over 2000 nanopores, but only 512 are monitored simultaneously. During the first few experiments on a flow cell, nanopores become inactive, but they are replaced with new working ones. Therefore, the first 4 to 5 experiments are practically done with the same number of pores. After that the number of working nanopores decreases by 5 to 10% per hour. It is worth noting that while most of the working nanopores exhibited comparable number of events within a factor of 5 from lowest to highest, a small percentage deviated and recorded markedly higher count of events. We attribute the occurrence of outliers to sample inhomogeneity within the microfluidics of a flow cell and/or to plausible differences within the same batch of protein nanopores, making a small percent of them highly conducive to translocation events. We call these pores “outliers” and found an approximate 2.5% of nanopores to be outliers. The analysis of the data we presented here includes the recordings from all channels including the recordings from the outliers. We performed an additional analysis of all the experiments by excluding the outliers, and even though the actual counts were less, the trends and conclusions are identical to the ones presented here.

### Event detection algorithm - OsBp_detect

Let *y* be an ordered sequence of real values representing a typical time-series obtained from a single Nanopore. We categorize regions of *y* into one of two states: current from an open channel, *I*_*o*_, where the Nanopore is unoccupied, and the residual current when some translocation event is taking place, *I*_*r*_. For the high-throughput characterization of a single translocation event corresponding to an OsBp-tagged oligonucleotide, we propose a segmentation algorithm that can determine the start and end positions in *y* for all events of interest based on user-defined thresholds. The analysis pipeline is divided into three steps:

1. Baseline current estimation
2. Identification of potential candidate events
3. Event filtering based on event features

First, the baseline *I*_*o*_, *b*_*o*_, is established by taking a median of the signal values between estimated lower and upper bounds, {*o*_*low*_, *o*_*up*_}, of the open current: *b*_*o*_ = med({*i*: *i* ∈ *y, o*_*low*_ < *i* < *o*_*up*_}). While the signal noise is dependent on the Nanopore platform used, the sharp transitions between the *I*_*o*_ and *I*_*r*_ states from osmium-tagged oligos permit the use of a single threshold-based parser for event detection pipeline. With this approach, events are identified if they pass a set threshold away from the local baseline level. The threshold is defined by parameter *b*_*all*_ which should be low enough to capture as many translocation events as possible (Fig. 8). By default, 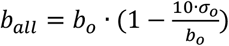 where σ_*o*_ represents the signal noise of the open current. The noise constant, σ_*o*_, is determined by splitting *y* into small segments (segment size used, *n* = 100,000) and calculating the global standard deviation of the open current signal values, σ({*i*: *i* ∈ *y, o*_*low*_ < *i* < *o*_*up*_}). The σ_*o*_ value additionally provides a useful quality control metric to detect pores with unstable baselines and large events rates or pores that have been blocked.

**Fig. 8:**
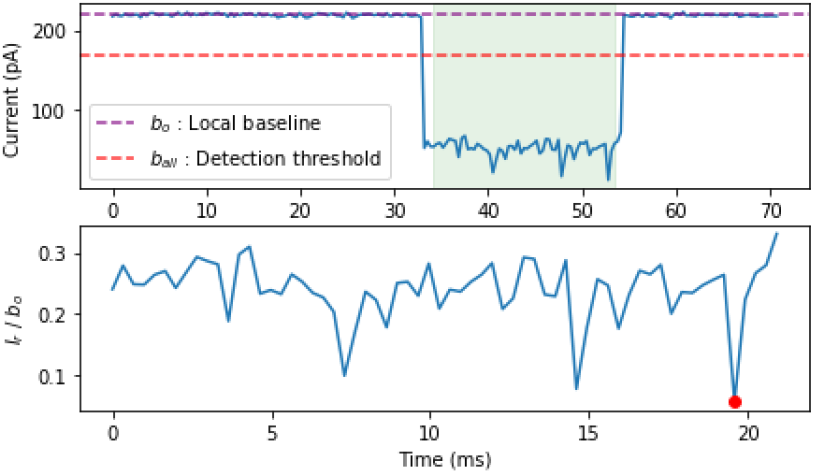
Stages of the event detection pipeline. *Top*: Following the estimation of the baseline open current, *b*_*o*_ outlined in purple, candidate events that cross the detection threshold, *b*_*all*_ outlined in purple, are identified and highlighted in green. *Bottom*: The identified event is accepted if it satisfies the event duration and min (*I*_*r*_/*b*_*o*_) thresholds. The min (*I*_*r*_/*b*_*o*_) value is highlighted in red.

Finally, for the qualification of valid translocation events, we apply two filtering conditions to the identified events. The filters correspond to the minimum and maximum event length defined by parameters, {*t*_*min*_, *t*_*max*_}, and the range of lowest residual current, {*b*_*min*_, *b*_*max*_}, expressed as a ratio with respect to *b*_*o*_. The event length thresholds, {*t*_*min*_, *t*_*max*_}, can be adjusted to monitor the speed of translocation whereas {*b*_*min*_, *b*_*max*_} enables the separation of OsBp-tagged and untagged oligo species. Let *t*_1_ and *t*_2_ represent the starting and ending indices for any given event in *y*. For 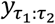 to be classified as a potential OsBp translocation event, both of the following conditions must hold:

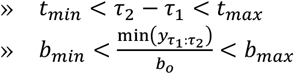

The event detection pipeline is available as a Python library, ‘*osbp_detect*’. A cross-platform graphical user interface has been included, to enable direct reporting of translocation events from ONT bulk *fast5* files (https://github.com/kangaroo96/osbp_detect).

## Data availability

All of the data generated during this study are included in this published article (and its Supplementary Information files). All of raw *i-t* data (each file is 2 to 6GB) may be obtained by request from AK.

**Supplementary Information**, see separate file.

## Acknowledgements

We are grateful to the SBIR/NIH program for their continued support. The work, JGB, ASWK, and AK were supported by SBIR grant 1R43HG010841 from NIH/NHGRI.

## Author Contributions Statement

A.K. conceived the project, designed the study and wrote the manuscript; she conducted the nanopore, and HPLC experiments and analyzed the data. W.J. designed and conducted the enzymatic polymerization reactions. A.S.W.K. developed the *OsBp_detect* software tool. J.G.B. helped with data analysis and manuscript preparation. All four authors reviewed the manuscript.

---

**Table :**
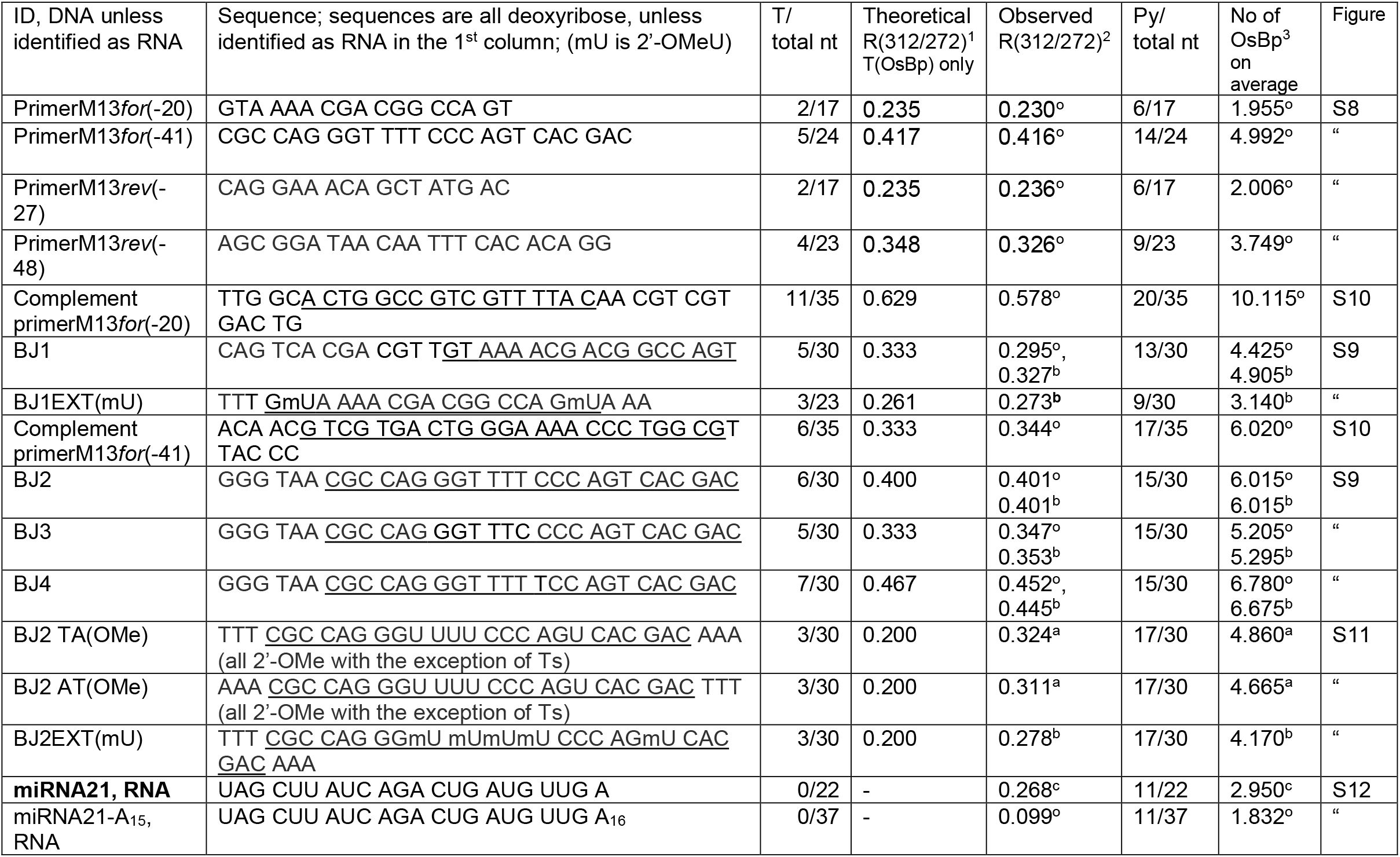
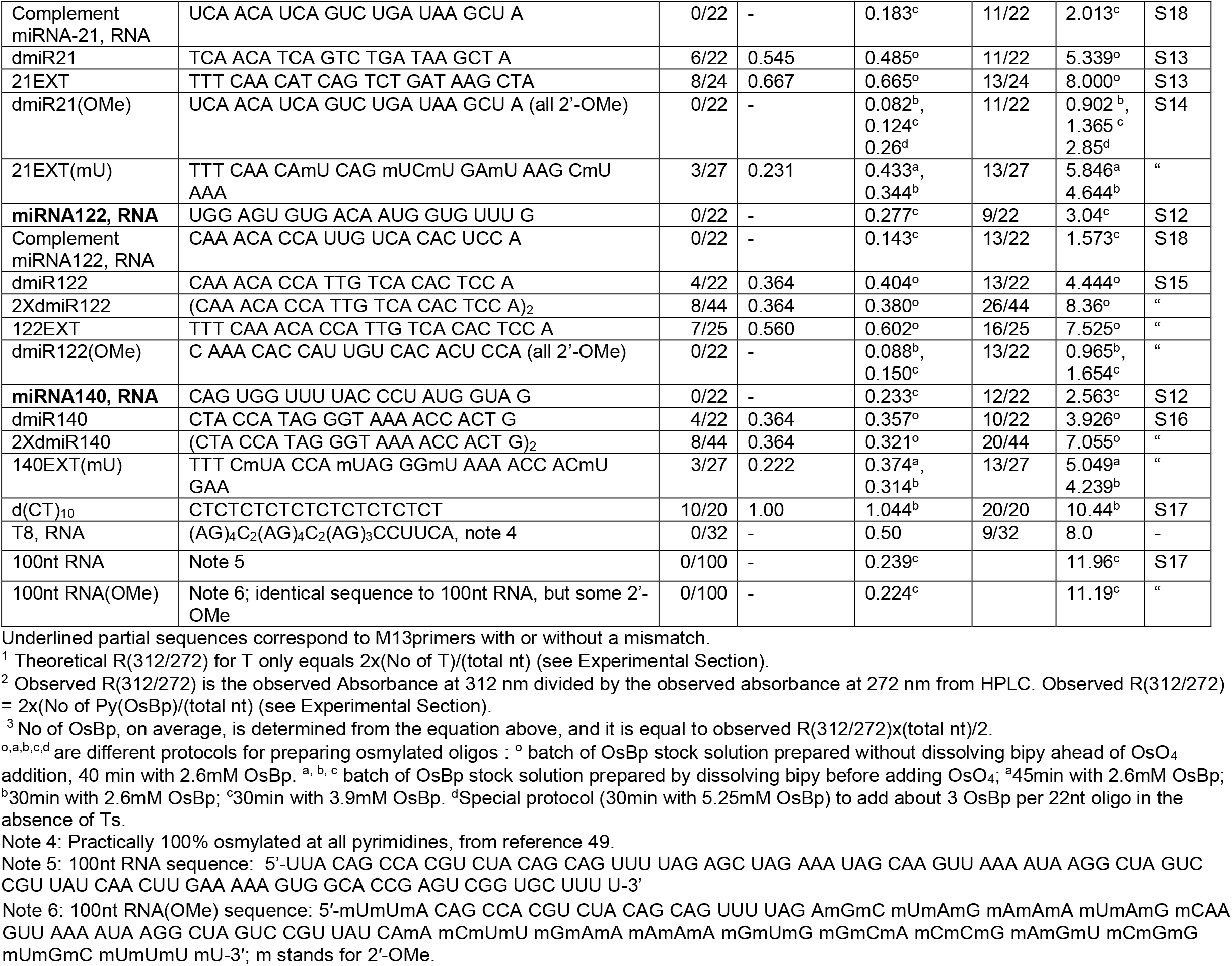
List of tested DNA or RNA oligos. The 3 miRNAs tested here, in bold, have the sequence of the corresponding miRNA-5p. Oligo ID, sequence, number of thymidines over total nucleotides (T/total nt), number of pyrimidines over total nucleotides (Py/nt), theoretical R(312/272) for T(OsBp) (see footnote and Experimental Section), Observed R(312/272) is the ratio of the observed areas under the HPLC peak at 312nm over the area at 272nm; HPLC analytical profiles for each oligo and its osmylated conjugate (Fig. S#) are included in the Supplementary Information. The number of Py(OsBp) or OsBp moieties, on average, depends on the protocol used for osmylation and is calculated as described in footnote 3.

