## SUPPLEMENTAL INFORMATION for "Ready-to-use nanopore platform for the detection of any DNA/RNA oligo at attomole range using an Osmium tagged complementary probe"

```
 _ _ _ _ _ _ _ _ _ _ _
 /\/\/\/\/\/\/\/\/\/\/\
( O | s | B | p | D | e | t | e | c | t )
 \_/\_/\_/\_/\_/\_/\_/\_/\_/\_/
============ Thresholds ============
Event duration (in tps): 4 - 300
Lowest Ir/Io < 0.55
All Ir/Io < 0.6
======================================
Processing channel 1...
400 events detected.
# Channel 1, Sampling rate: 3012.0 Hz, Io: 251.7634153366089 pA
```

| | | | |
|---|---|---|---|
| 1 | 353587 | 353611 | 0.06775701 |
| 2 | 398370 | 398375 | 0.11623832 |
| 3 | 407336 | 407348 | 0.08820094 |
| 4 | 412256 | 412438 | 0.21670561 |
| 5 | 447187 | 447197 | 0.07593458 |
| 6 | 453217 | 453249 | 0.06600467 |
| 7 | 492322 | 492371 | 0.07009346 |
| 8 | 493357 | 493361 | 0.32476636 |
| 9 | 497867 | 498043 | 0.21728972 |
| 10 | 520164 | 520179 | 0.19742991 |
| 11 | 542446 | 542455 | 0.06074766 |
| 12 | 567601 | 567612 | 0.08528037 |
| 13 | 595027 | 595031 | 0.33294393 |
| 14 | 650430 | 650488 | 0.08002336 |
| 15 | 650828 | 650858 | 0.16647196 |
| 16 | 742841 | 743058 | 0.04614486 |
| 17 | 792186 | 792318 | 0.12733645 |
| 18 | 832875 | 832913 | 0.04672897 |
| 19 | 841164 | 841188 | 0.28387851 |
| 20 | 907859 | 907868 | 0.27686916 |
| 21 | 1073757 | 1073771 | 0.19801402 |
| 22 | 1075537 | 1075541 | 0.34521028 |
| 23 | 1080276 | 1080460 | 0.15887851 |
| 24 | 1081066 | 1081078 | 0.15478972 |
| 25 | 1101348 | 1101364 | 0.21378505 |
| 26 | 1110665 | 1110672 | 0.26109813 |
| 27 | 1132460 | 1132678 | 0.23948598 |
| 28 | 1178961 | 1179019 | 0.39252336 |
| 29 | 1211889 | 1211901 | 0.26518692 |
| 30 | 1267382 | 1267386 | 0.28679907 |

```
 _ _ _ _ _ _ _ _ _ _ _
 /\/\/\/\/\/\/\/\/\/\/\
( O | s | B | p | D | e | t | e | c | t )
 \_/\_/\_/\_/\_/\_/\_/\_/\_/\_/
============ Thresholds ============
Event duration (in tps): 4 - 300
Lowest Ir/Io < 0.55
All Ir/Io < 0.6
======================================
Processing channel 1...
Processing channel 2...
Processing channel 3...
Processing channel 4...
39 events detected.
# Channel 4, Sampling rate: 3012.0 Hz, Io: 240.23457124829292 p
```

| | | | |
|---|---|---|---|
| 1 | 567674 | 567705 | 0.09669523 |
| 2 | 766999 | 767209 | 0.29681763 |
| 3 | 918896 | 918925 | 0.09669523 |
| 4 | 1050173 | 1050204 | 0.30966952 |
| 5 | 1599517 | 1599525 | 0.20379437 |
| 6 | 1680746 | 1680758 | 0.28580171 |
| 7 | 2049151 | 2049162 | 0.16034272 |
| 8 | 2051214 | 2051247 | 0.18665851 |
| 9 | 2087063 | 2087068 | 0.18359853 |
| 10 | 2297779 | 2297785 | 0.15422277 |
| 11 | 2544443 | 2544458 | 0.26376989 |
| 12 | 2862924 | 2862940 | 0.28335373 |
| 13 | 3044089 | 3044100 | 0.41003672 |
| 14 | 3150965 | 3150971 | 0.35495716 |
| 15 | 4810901 | 4810911 | 0.41554468 |
| 16 | 4810943 | 4810950 | 0.41615667 |
| 17 | 4811778 | 4811783 | 0.48347613 |
| 18 | 4812081 | 4812139 | 0.41432069 |
| 19 | 4812197 | 4812203 | 0.46511628 |
| 20 | 5915973 | 5915996 | 0.43329253 |
| 21 | 5916909 | 5916964 | 0.41432069 |
| 22 | 6204350 | 6204361 | 0.29620563 |
| 23 | 6218284 | 6218451 | 0.13219094 |
| 24 | 6314244 | 6314248 | 0.33659731 |
| 25 | 6533152 | 6533163 | 0.24235006 |
| 26 | 6694009 | 6694020 | 0.34210526 |
| 27 | 6716411 | 6716419 | 0.19828641 |
| 28 | 7095854 | 7095865 | 0.43941249 |
| 29 | 7136165 | 7136282 | 0.45777234 |
| 30 | 7214510 | 7214528 | 0.18849449 |

**Figure S1: Samples of *tsv* files, obtained by running the *OsBp_detect* software on *fast-5* files.** Left, sample probe T8(RNA); right, sample is a mixture of $d(CT)_{10}$:T8(RNA)=1:1, both in about 90% ONT buffer (see caption of Fig. 2a for experimental conditions).

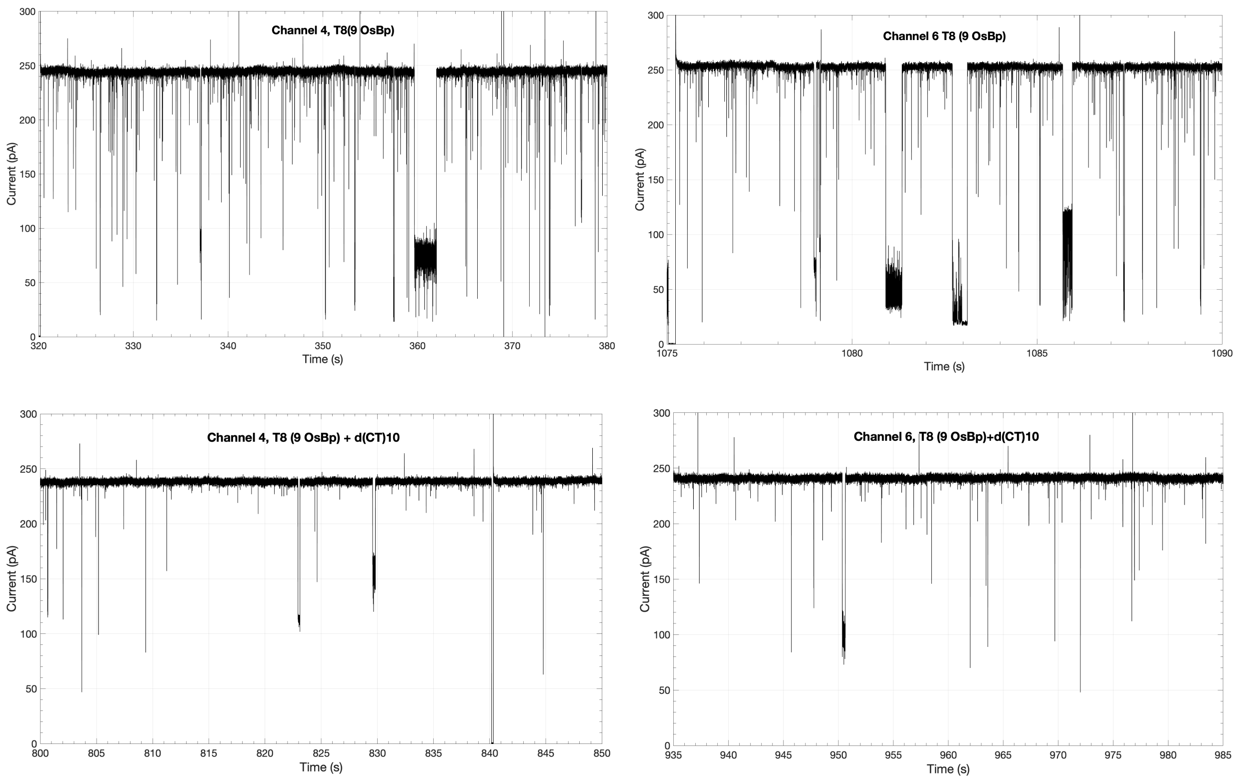


**Figure S2: *I-t* recordings from two nanopore experiments ranging from 15 to 60s.** Top, probe T8(RNA), *i-t* recordings from two different channels. Bottom, mixture of $d(CT)_{10}$ : T8(RNA)=1:1, *i-t* recordings from the same two channels as on the top. Vertical lines that cross the x-axis (=0 pA) are instrument generated lines by voltage reversal, and not events. Top recordings show multiple and deep events, bottom recordings show few and shallow events. Shallow events are attributed to molecules bumping at the pore aperture, without traversing the pore and they are not counted when selecting “All Ir/Io < 0.6” (see Fig. S1, and Experimental Section).

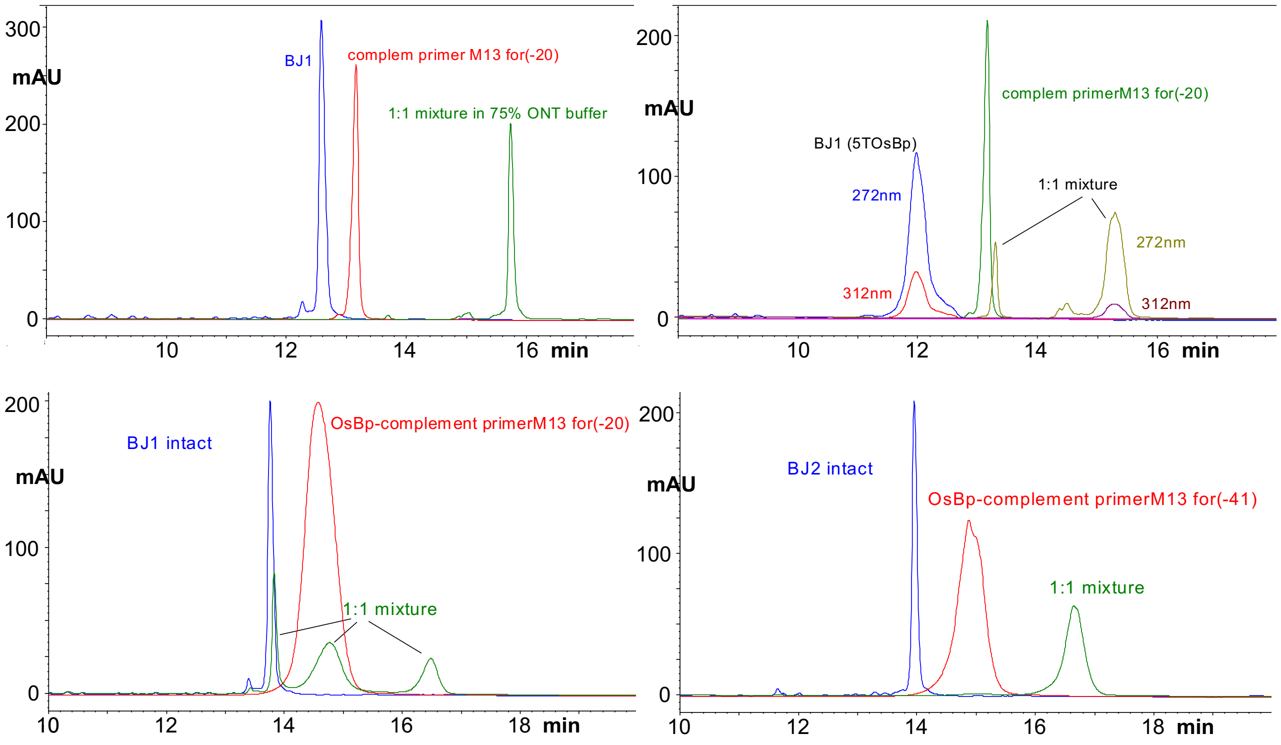


**Figure S3: HPLC profiles of individual components and their 1:1 mixtures in a sample solvent about 90% ONT buffer.** All HPLC profiles shown at 260 nm, with the exception of the osmylated probe and hybrid in the top right profile that is shown at 272 nm and 312 nm. Hybridization is evident by a mixture HPLC profile where the main peak elutes after the individual components. Top left, hybridization shown for two intact oligos, BJ1 and its complementary intact oligo (complement primerM13*for*(-20)). See Fig. 3b for comparable result with another pair of intact oligos. Top right, hybridization shown for probe BJ1 with 5 OsBp out of 30nt, and its complementary intact oligo (complement primerM13*for*(-20)). Left bottom, only partial hybridization is shown for intact BJ1 and its complementary osmylated complement primerM13*for*(-20) with 11 OsBp out of 35nt. Right bottom, hybridization is shown for intact BJ2 and its complementary osmylated complement primerM13*for*(-41) with 6 OsBp out of 35nt. HPLC method B is used for analysis (see Experimental Section).

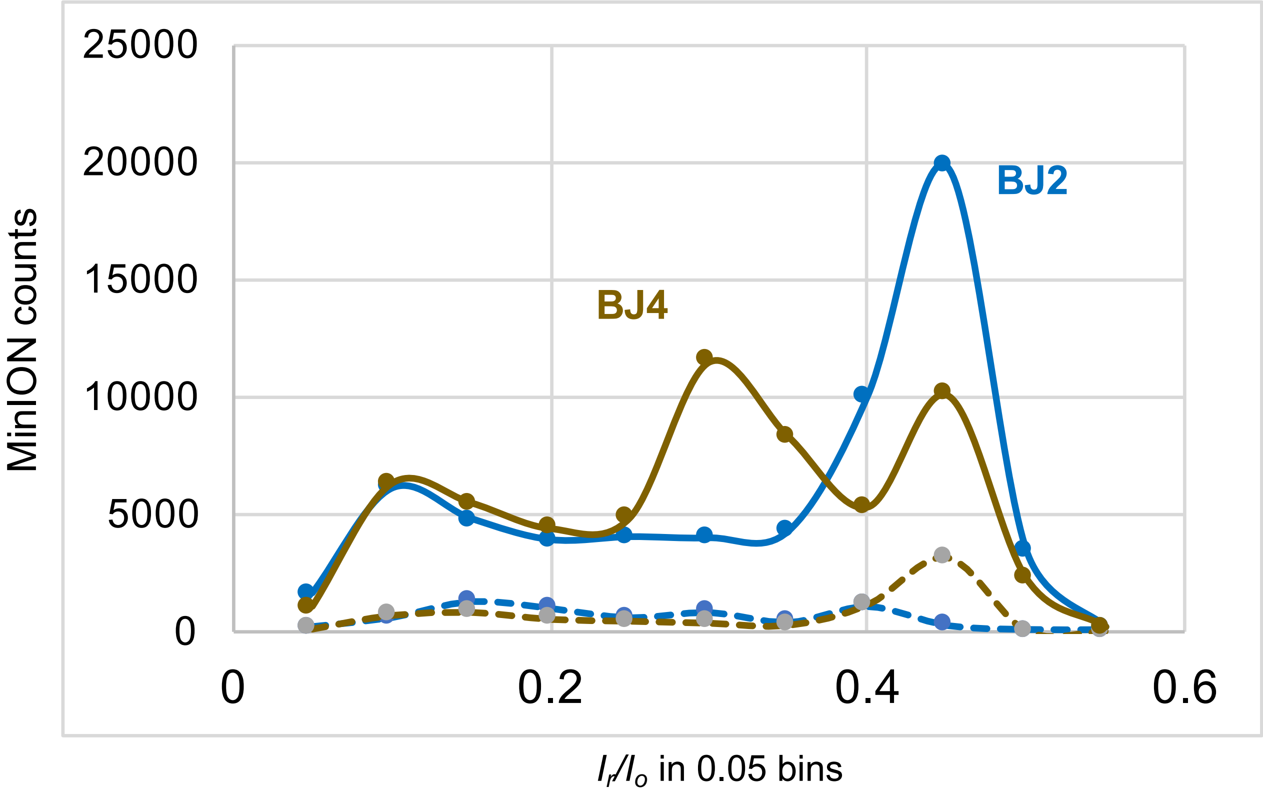


**Figure S4**: **Nanopore experiments with probes BJ2 and BJ4 show few events at -180mV and numerous events at -220mV.** Probes BJ2, blue traces, and BJ4, brown traces, tested at -180mV (dashed traces) show negligible number of counts. No new sample was added, the voltage was raised to -220mV and probes were tested at -220 mV (solid traces) and numerous events were detected**.** Both probe samples were used at a 0.2nmole load. Experiments were conducted on the same flow cell in the order BJ2 at -180mV, BJ2 at -220mV, BJ4 at -180mV and BJ4 at -220mV; the duration of each experiment was 1h. Data acquisition and analysis as described under Fig. 2a (see text).The dramatic difference in counts at the different applied voltage clearly suggests that these probes and other probes, of similar design, do not traverse the proprietary CsGg nanopore at the lower voltage and require high applied voltage of about -220mV to translocate.

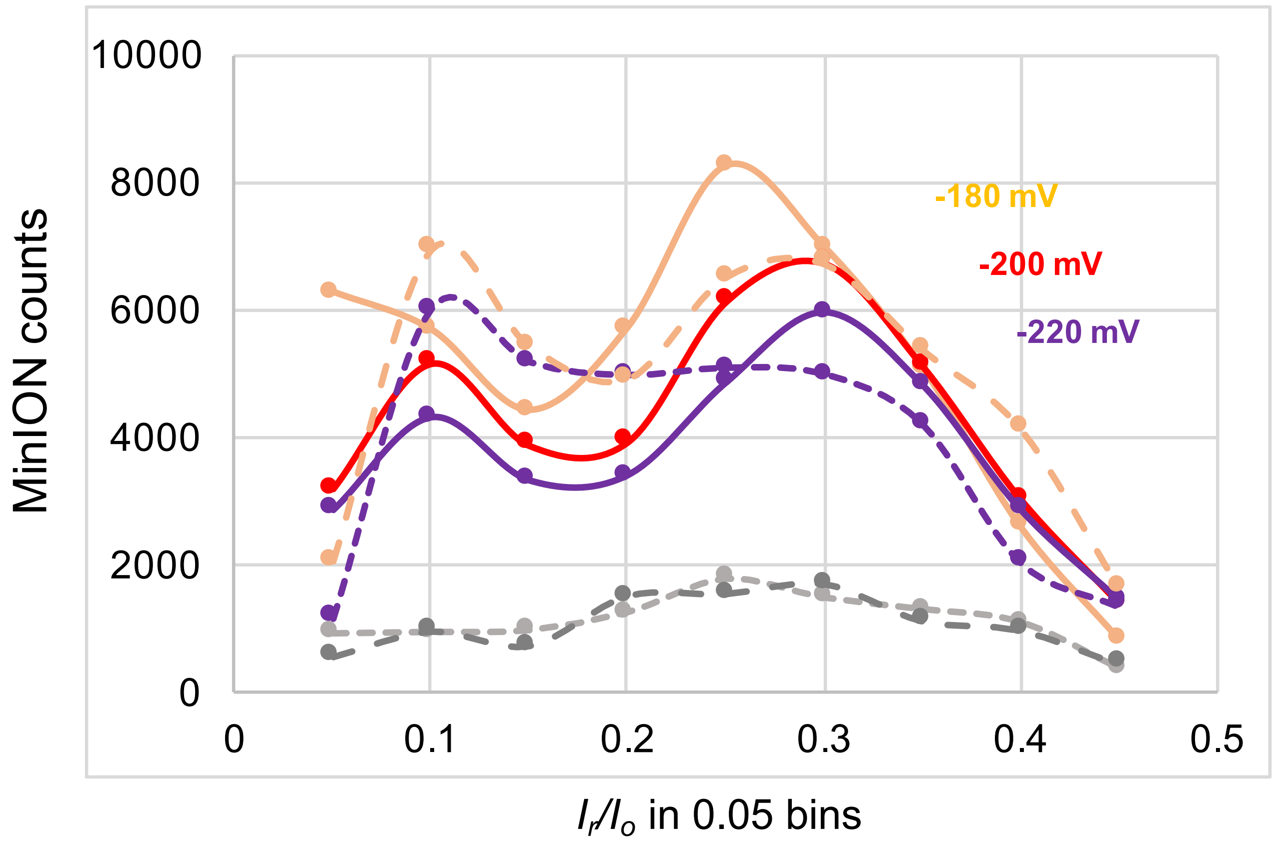


**Figure S5: Effect of applied voltage on the counts observed with a mixture of intact miRNA122 and miRNA140** at 10μM each at -180mV (orange trace), at -200mV (red trace), at -220mV (purple trace). Data shown at -180mV (dashed orange trace), are the same data as in 2b (Flongle, red trace)) but normalized by multiplying with 10, as the MinION has about 10 times more working channels compared to the Flongles that we purchased/used. Increased applied voltage reduces slightly the count of events, consistent with faster translocation and reduced detectability. An experiment with 10μM of miRNA21-$A_{15}$ at -220mV (dashed purple line) exhibits comparable counts with the combo of miRNA122 and miRNA140, but a distinct profile compared to the miRNA with no $A_{15}$-tail. The effect of voltage on the intact RNAs is in stark contrast to the effect of voltage on most of the probes tested in this study. No detectable difference in counts is observed with the control/buffer between -200 and -220mV (light and dark dashed grey traces, respectively).

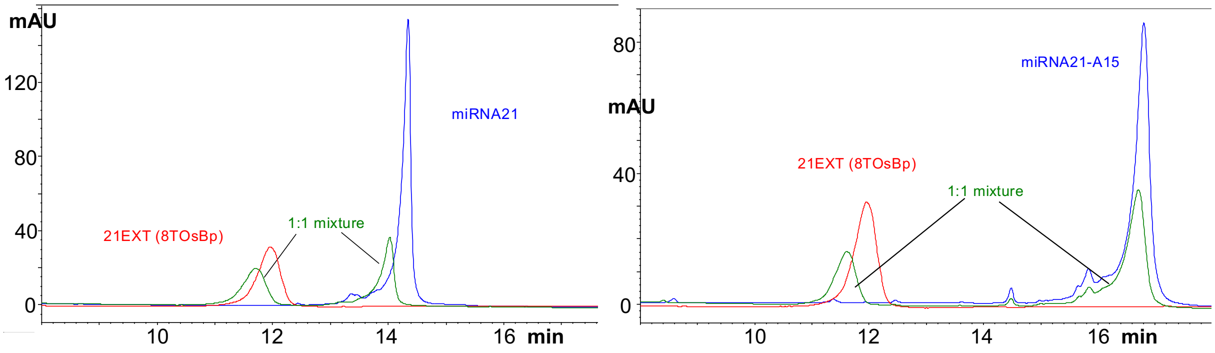


**Figure S6: HPLC profiles of 1:1 mixtures (non-hybrids) of miRNA21 or miRNA21-A15 using probe 21EXT (with 8 T(OsBp),** see sequences in table). Left profile the same as in Figure 3c, repeated here for direct comparison with the right profile. The difference in these two cases is that, due to the added $A_{15}$-tail, miRNA21-$A_{15}$ elutes couple of min later compared to miRNA21.
Samples in about 90% ONT buffer as the sample solvent. HPLC profiles obtained with HPLC method B (Experimental Section). HPLC profile of the mixture sample matches closely the sum of the HPLC profiles of the two components, providing evidence for no detectable hybridization in these two cases.

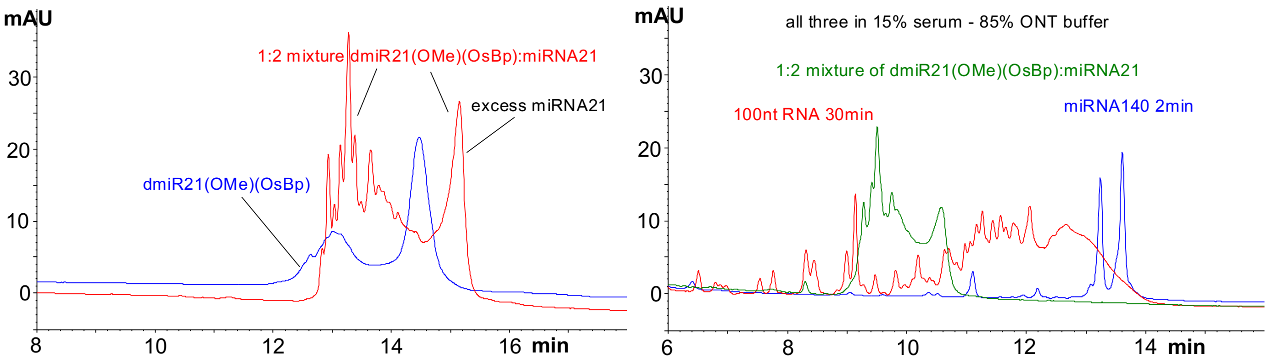


**Figure S7: Stability of hybrid in 15% serum -85% ONT buffer.** Left, repeat of Fig. 7a (red trace) in order to compare directly with HPLC profile (green trace) to the right. Right: HPLC profiles of three samples in 15% serum -85% ONT buffer: miRNA140 (blue trace) 2min incubation before analysis, 100nt RNA (red trace) 30min incubation before analysis. The longer incubation is why the degradation of 100nt RNA appears more severe compared to the degradation of miRNA140. Mixture of dmiR21(OMe)(OsBp): miRNA21=1:2 in about 5% water- 95% ONT buffer (left, red trace) and the same mixture in 15% serum -85% ONT buffer (right, green trace). The HPLC profiles (red trace in left profile and green trace in right profile) appear comparable suggesting that the hybrid suffers insignificant degradation in 15% serum - 85% ONT buffer. HPLC method B used for these analyses (see Fig. S10 and Experimental Section).

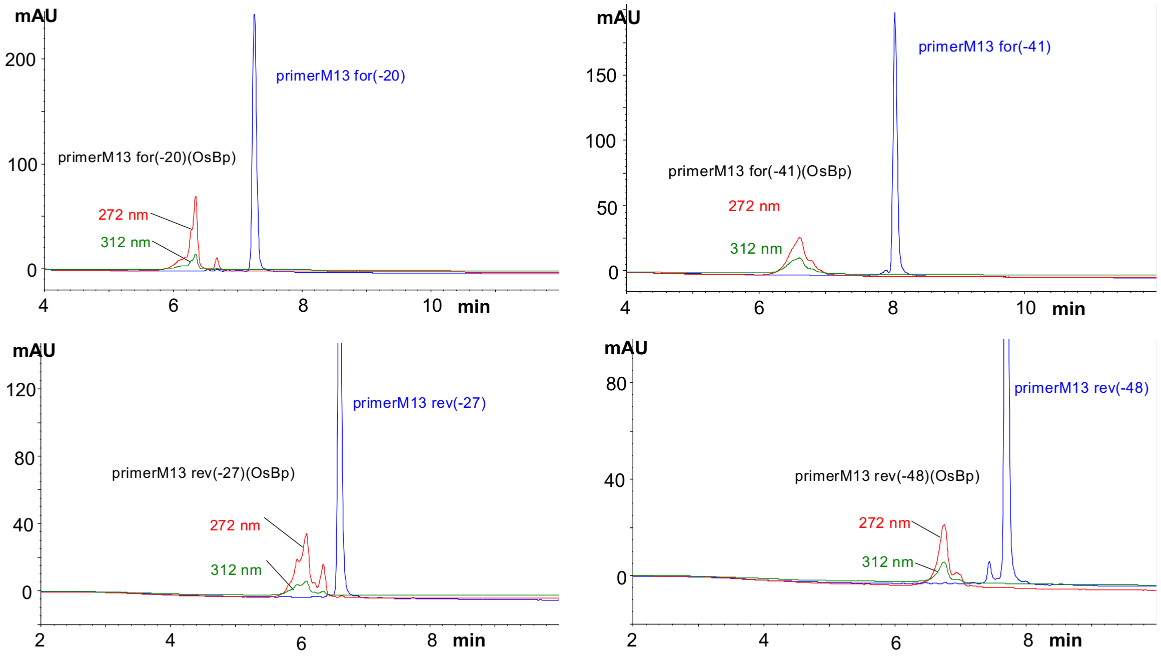


**Figure S8: HPLC profiles of the four intact M13 primers and their corresponding T-osmylated derivatives,** using HPLC method A (Experimental Section). Analysis of oligos in water as the sample solvent. T-osmylation of these oligos was conducted using protocol o (see Experimental Section). The reason osmylated oligos appear as multiple peaks is because top or bottom addition of OsBp to the C5-C6 double bond leads to topoisomers (see text), that this chromatography resolves.

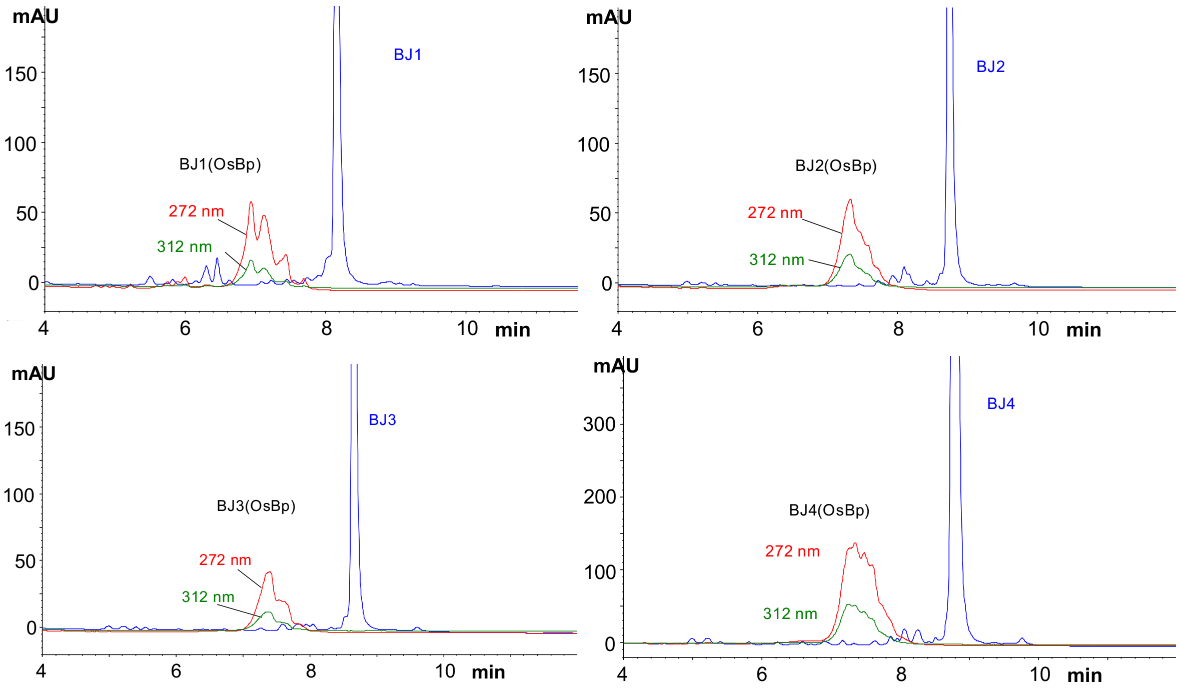


**Figure S9: HPLC profiles of the four intact BJ1-4 and their corresponding T-osmylated derivatives** using HPLC method A. Analysis of oligos in water as the sample solvent. T-osmylation of these oligos was conducted using protocol o (see Experimental Section).

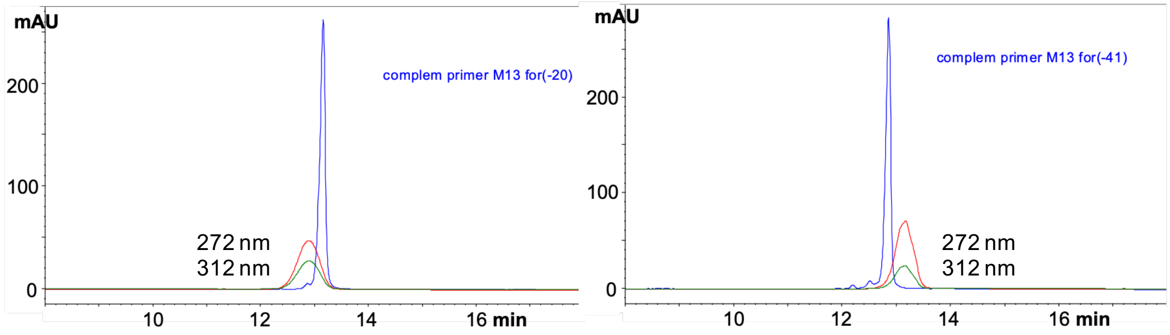


**Figure S10: HPLC profiles of the 2 intact complements of primerM13*for*(-20) and primerM13*for*(-41) shown at 260 nm, and their corresponding T-osmylated derivatives** shown at 272nm and 312nm using HPLC method B (see Experimental Section). Analysis of oligos in water as the sample solvent. Briefly HPLC Method B is using the DNA Pac PA200 HPLC column from ThermoFisher Scientific at the 2x250mm configuration with 0.45mL/min flow and 15ºC column compartment. Solvents are aqueous pH 8.0±0.2 mobile phases A (MPA) and mobile phase B (MPB) with 25mM TRIS.HCL buffer; MPB is 1.5 M NaCl. Initial conditions are 90% MPA – 10% MPB, and the gradient is from 10% to 50% MPB in 20 min. The total analysis time including column equilibration is 30min. T-osmylation of these oligos was conducted using protocol o (see Experimental Section). Right profile shows an atypical, but confirmed result, namely an osmylated conjugate that elutes later compared to the parent intact nucleic acid.

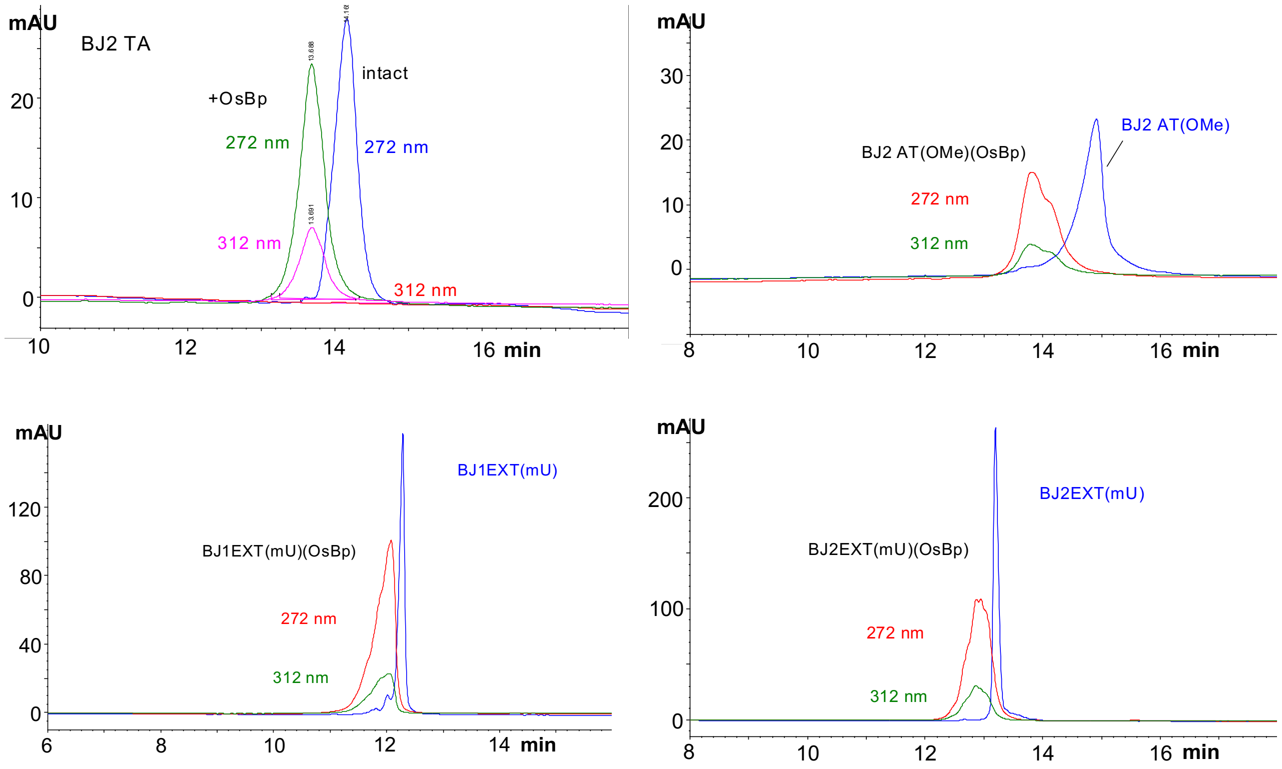


**Figure S11: HPLC profiles of advanced probe designs and their osmylated derivatives.** BJ2 TA(OMe) (top, left) and BJ2 AT(OMe) (top, right), as well as BJ1EXT(mU) (bottom, left) and BJ2EXT(mU) (bottom, right) shown at 260nm and their corresponding T-osmylated derivatives shown at 272nm and 312nm (see sequences in table in text). HPLC profiles obtained with HPLC method B (see Fig. S10, and Experimental Section). Materials in water as sample solvent. A nanopore experiment conducted with probe BJ2 TA(OMe) indicated excellent translocation properties with numerous counts at a relatively low probe load (Fig. 4d). A nanopore experiment conducted with BJ2 AT(OMe) at comparable conditions was not conclusive. Since the only difference between the two probes is that the 3'-end on one is the 5'-end on the other, it is not clear whether or not BJ2 TA(OMe) with the Ts at the 5'-end and the As at the 3'-end is a superior probe compared to BJ2 AT(OMe) with the Ts at the 3'-end and the As at the 5'-end.

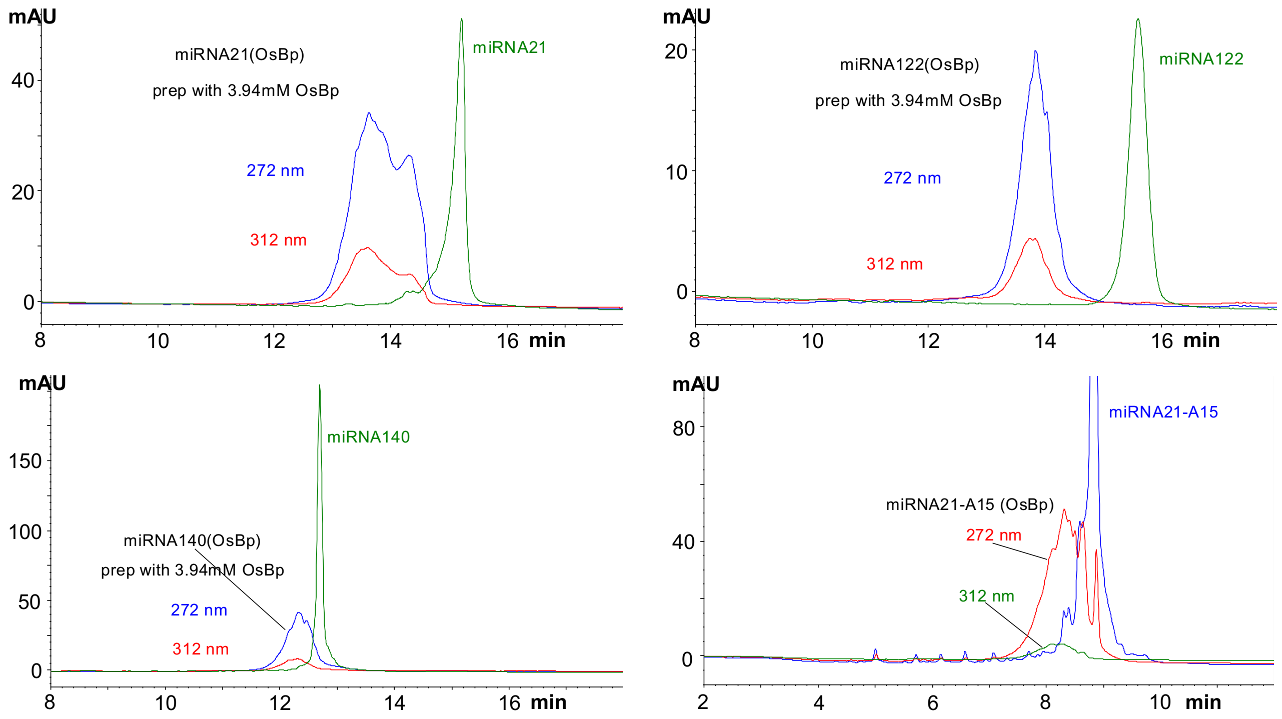


**Figure S12: HPLC profiles of the intact miRNA21, miRNA122, miRNA140 and miRNA21-A$_{15}$ (these are the -5p sequences) shown at 260nm and their corresponding partially osmylated derivatives;** different materials are analyzed at different sample load. Osmylation protocol o was used for miRNA21-A$_{15}$ and protocol c for the other 3 miRNAs (see table and Experimental Section). HPLC method A used for the analysis of miRNA21-A15 and HPLC method B for analysis of the other 3 miRNAs (see Experimental Section). Materials in water as the sample solvent.

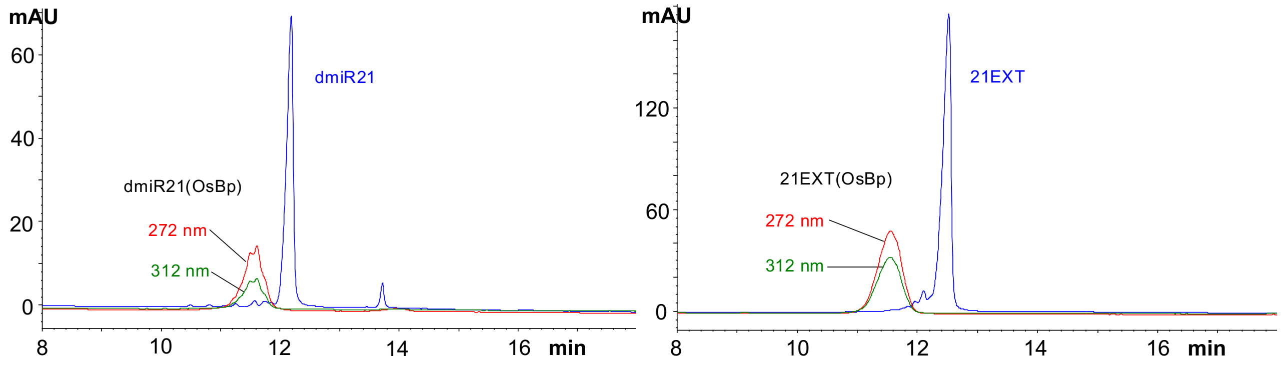


**Figure S13: HPLC profiles of unsuccessful probes for miRNA21.** Left, HPLC profiles of intact dmiR21 at 260nm and the T-osmylation product at 272nm and 312nm. Right, HPLC profiles of intact 21EXT and its T-osmylation product. Osmylation was carried out using protocol o 40min with 2.63mM OsBp, earlier process where bipy was dissolved after adding $OsO_4$ (see Experimental Section). Materials in water for analysis, and analysis was done using HPLC method B (see Fig. S10 and Experimental Section).

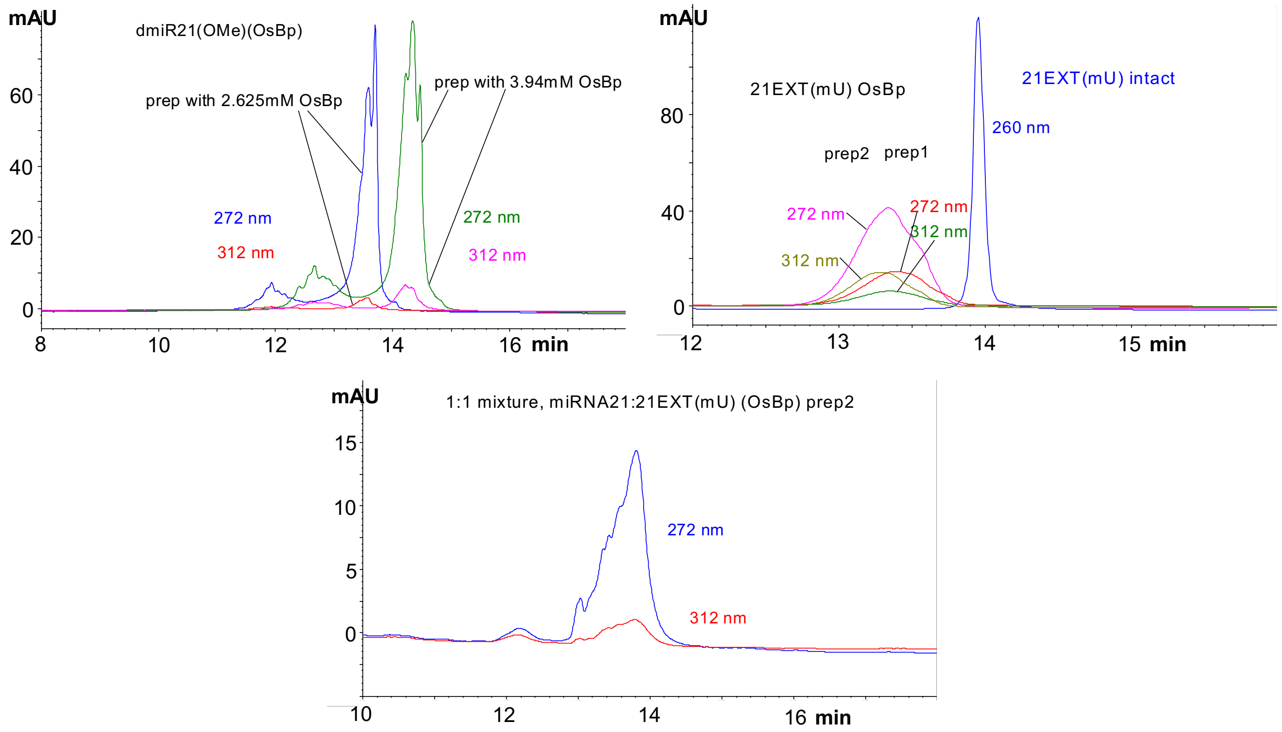


**Figure S14**: **HPLC profiles of successful probes for miRNA21.** Left, HPLC profiles of the osmylation products of dmiR21(OMe) using protocol b 30min with 2.63mM OsBp and protocol c 30min with 3.94mM OsBp. A third protocol (d) with 30min incubation using 5.25mM OsBp was used to osmylate this probe for the nanopore experiment (Fig. 6b). This was necessary because dmiR21(OMe) contains no Ts, and using protocols b and c results in very low levels of osmylation and diminished detectability. Right, probe 21EXT(mU) intact and its osmylation products with the two different protocols; prep1, using protocol a 45min with 2.63mM OsBp and prep2 protocol b 30min with 2.63mM OsBp. Bottom profile, repeat of Fig. 7c in order to compare the HPLC profile of the hybrid with the HPLC profile of the probe alone (b, above) and see that they are distinct. Analysis was done using HPLC method B (see Fig. S10 and Experimental Section).

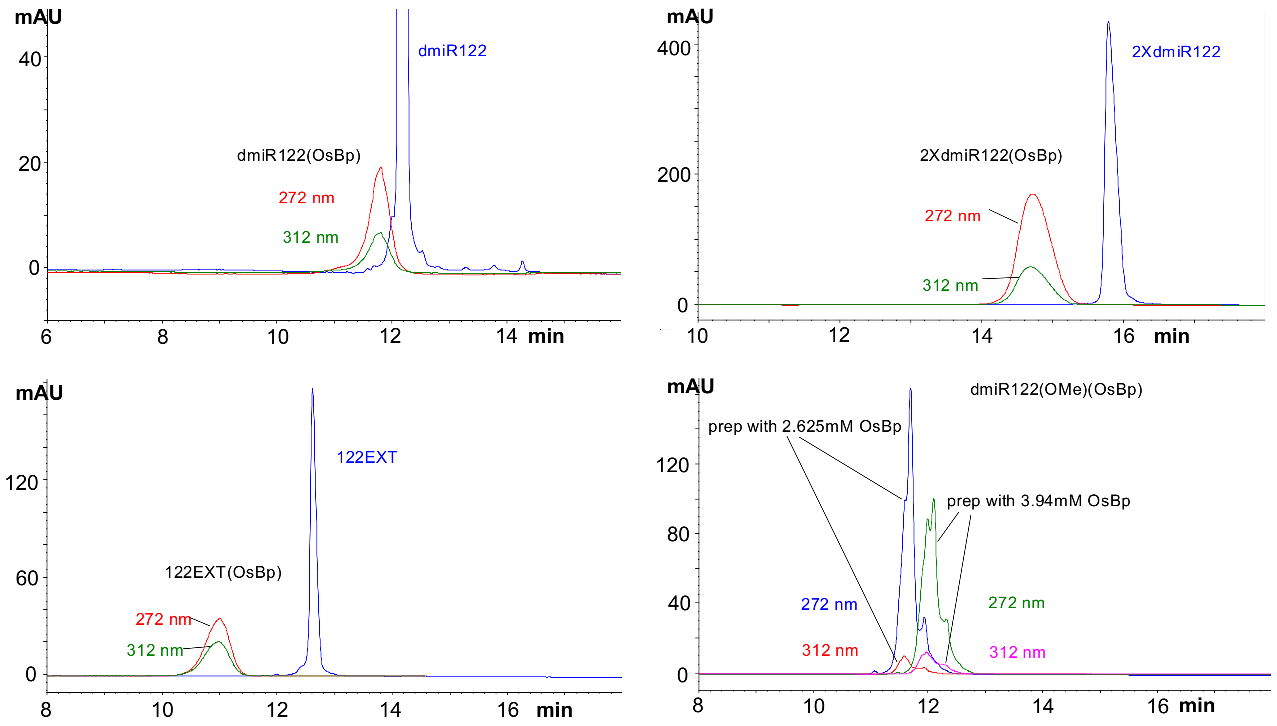


**Figure S15: HPLC profiles of successful probes for miRNA122.** HPLC profiles of the intact miRNA122 probes shown at 260nm and their corresponding T-osmylated derivatives (the actual probes) at 272 and 312nm. Osmylation protocol o was used to osmylate dmiR122, 2XdmiR122 and 122EXT. dmiR122(OMe) does not have any T, and was osmylated using protocols b or c (see table for sequences and for protocols). Materials were in water for analysis and analysis was done using HPLC method B (see Fig. S10 and Experimental Section).

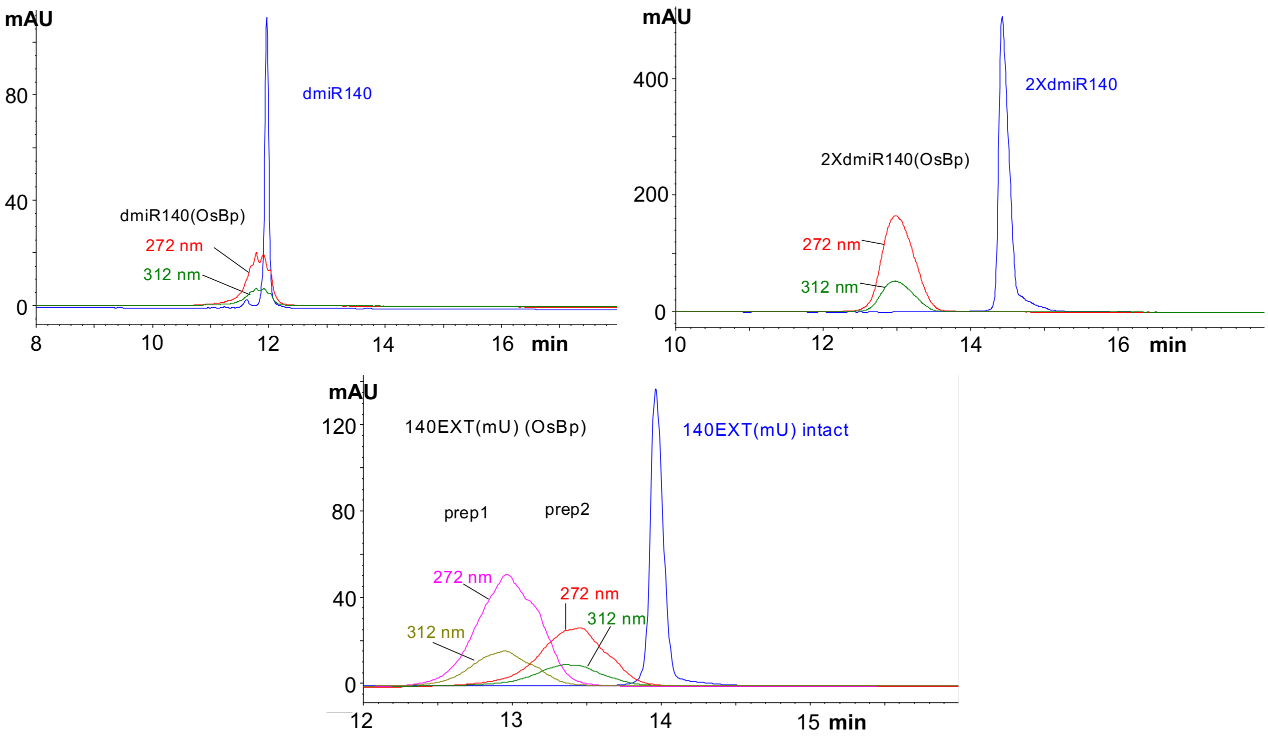


**Figure S16: HPLC profiles of the intact miRNA140 probes shown at 260nm and their corresponding osmylated derivatives** (the actual probes) shown at 272nm and 312nm. Top HPLC profiles with dmiR140 (left) and 2XdmiR140 (right) using osmylation protocol o, 40min with 2.63mM OsBp (earlier process, bipy not dissolved prior to $OsO_4$ addition. Bottom, probe 140EXT(mU) intact and its osmylation products with the two different protocols; prep2, using protocol a, 45min with 2.63mM OsBp, and prep1 protocol b, 30min with 2.63mM OsBp. Materials in water for analysis, and analysis was done using HPLC method B (see Fig. S10 and Experimental Section).

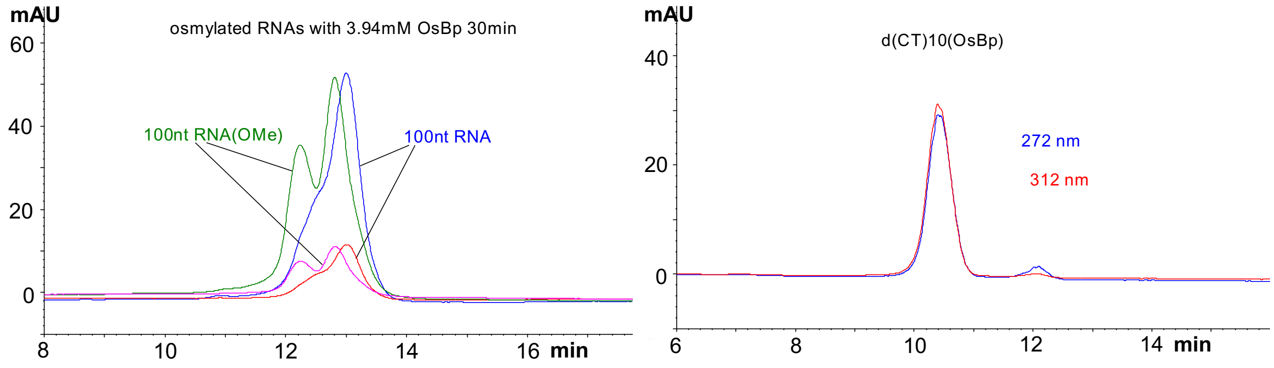


**Figure S17: HPLC profiles of partially osmylated RNAs and DNA.** Left, HPLC profiles of partially osmylated 100nt RNA and 100nt RNA(OMe) using osmylation protocol c (see table and Experimental Section). Right, HPLC profiles of T-osmylated $d(CT)_{10}$ using protocol b. Materials in water as the sample solvent. HPLC method B was used for all the samples and HPLC profiles are shown at 272nm and 312nm. Osmylation protocols and HPLC method can be found in the Experimental Section.

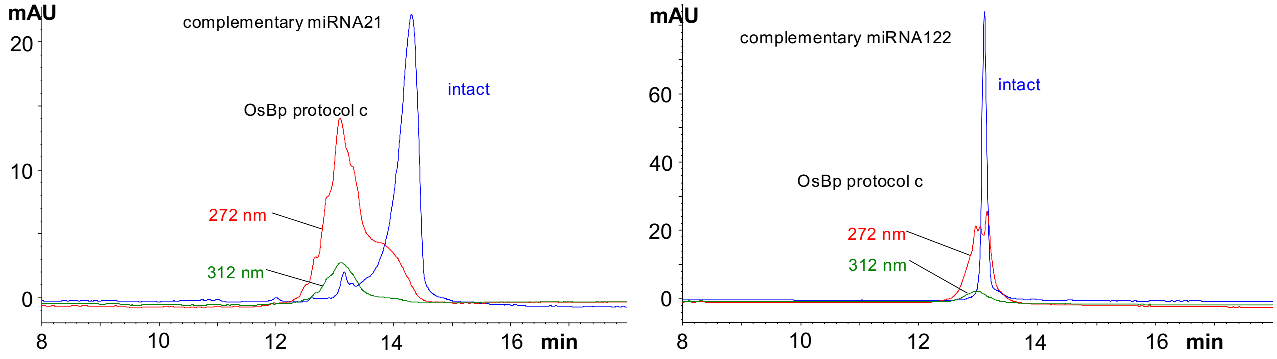


**Figure S18: HPLC profiles of partially osmylated 22nt RNAs.** Left, complementary miRNA21 and Right, complementary miRNA122. Osmylation protocol c was used, 30min with 3.94mM OsBp.

**Table S1:** Entries obtained from table in the text: Osmylation protocol c (30min with 3.94mM OsBp)

| Oligo ID | No of U in the sequence | No of Py(OsBp) for oligos with native U | No of Py(OsBp) for oligos with modified U | comment |
|---|---|---|---|---|
| miRNA21 | 8 | 2.950 | | |
| Complement of miRNA21, RNA | 6 | 2.013 | | |
| miRNA122 | 7 | 3.040 | | |
| dmiR21(OMe) | 6 | | 1.365 | All bases 2'-OMe |
| Complement of miRNA122, RNA | 4 | 1.573 | | |
| dmiR122(OMe) | 4 | | 1.65 | All bases 2'-OMe |
| 100nt RNA | 27 | 11.96 | | |
| 100nt RNA(OMe) | | | 11.19 | 50% of bases 2'-OMe |

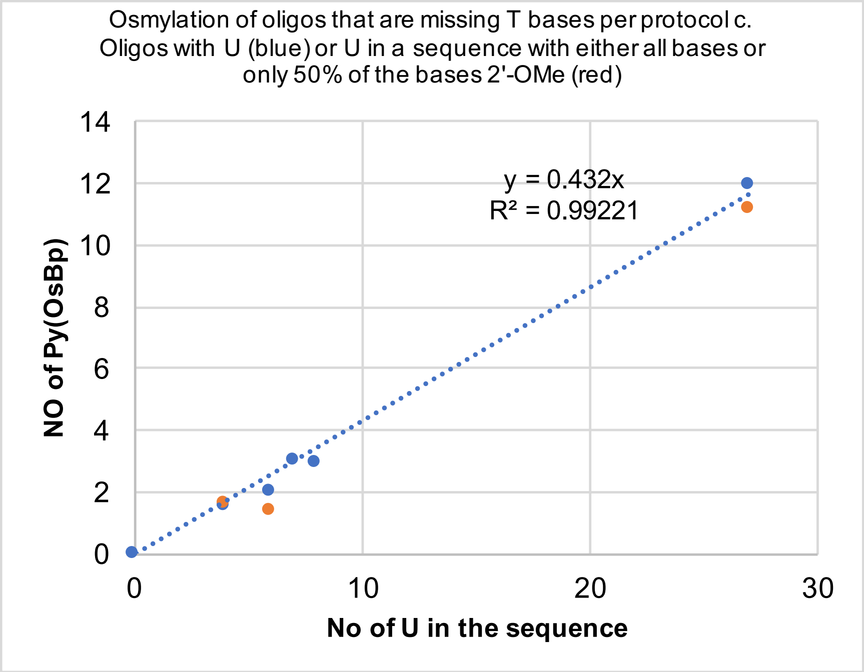


**Figure S19: Graph using the data from Table S1 above. A good linear correlation is obtained for the number of osmylated pyrimidines in an oligo, which is missing Ts, as a function of the number of U in a sequence.** The correlation does not appear to depend heavily on whether or not the sequence is a DNA, RNA or carries 2'-OMe groups on all or on a portion of the bases. The linear correlation may be used to estimate the number of osmylated pyrimidines per protocol c for any given sequence. The linear correlation is attributed to the observation that U is osmylated 4.7-times faster compared to C[44], and to the conditions of protocol c that yield only a small percentage of osmylated pyrimidines, and not a practically 100% osmylated oligo.
